# Reconstructing microvascular network skeletons from 3D images: what is the ground truth?

**DOI:** 10.1101/2024.02.01.578347

**Authors:** Claire Walsh, Maxime Berg, Hannah West, Natalie A. Holroyd, Simon Walker-Samuel, Rebecca J. Shipley

## Abstract

Structural changes to microvascular networks are increasingly highlighted as markers of pathogenesis in a wide range of disease, e.g. Alzheimer’s disease, vascular dementia and tumour growth. This has motivated the development of dedicated 3D imaging techniques, alongside the creation of computational modelling frameworks capable of using 3D reconstructed networks to simulate functional behaviours such as blood flow or transport processes. Extraction of 3D networks from imaging data broadly consists of two image processing steps: segmentation followed by skeletonisation. Much research effort has been devoted to segmentation field, and there are standard and widely-applied methodologies for creating and assessing gold standards or ground truths produced by manual annotation or automated algorithms.

The Skeletonisation field, however, lacks widely applied, simple to compute metrics for the validation or optimisation of the numerous algorithms that exist to extract skeletons from binary images. This is particularly problematic as 3D imaging datasets increase in size and visual inspection becomes an insufficient validation approach. In this work, we first demonstrate the extent of the problem by applying 4 widely-used skeletonisation algorithms to 3 different imaging datasets. In doing so we show significant variability between reconstructed skeletons of the same segmented imaging dataset. Moreover, we show that such a structural variability propagates to simulated metrics such as blood flow. To mitigate this variability we introduce a new, fast and easy to compute super-metric that compares the volume, connectivity, medialness, correct bifurcation point identification and homology of the reconstructed skeletons to the original segmented data. We then show that such a metric can be used to select the best performing skeletonisation algorithm for a given dataset, as well as to optimize its parameters. Finally, we demonstrate that the super-metric can also be used to quickly identify how a particular skeletonisation algorithm could be improved, becoming a powerful tool in understanding the complex implication of small structural changes in a network.

## 1. Introduction

Vascular networks are complex, interlinked, three-dimensional structures,which play a fundamental role in homeostasis and can be biomarkers of disease. A range of imaging techniques and image processing methods have been developed to image and quantitatively analyse them including magnetic resonance imaging (MRI), X-ray computed tomography (CT) and ultrasound (1; 2; 3) for larger vessel, and multi-photon microscopy, ultra fast ultrasound and photoacoustic imaging for the smaller microvasculature (3; 4). Recently, entire blood vessel networks in large tissue samples have been reconstructed using three-dimensional microscopy methods such as lightsheet microscopy, optical projection tomography (OPT), Multi-fluorescent high-resolution episcopic microscopy (MF-HREM) and Hierarchical Phase-Contrast Tomography (HiP-CT) (5; 6; 7; 8; 9). Each of these imaging techniques generates images with contrast of blood vessel location, alongside other structures and measurement noise. In order to utilise these data the vascular network must be digitized from the images requiring a two stage process: 1) Segmentation - distinguishing voxels within a blood vessel from the background or noise, 2) Skeletonisation - reducing the 3D voxel representation to a skeleton representation of segments, with each segment defined by start and end nodes, length, radius and the connections to other segments.

These image processing steps are pivotal to understanding the link between structure and function for microvascular networks in health and disease: a link that is often subtle and hard to fully characterise. For example, Alzheimer’s Disease (AD) is know to have associated vascular dysfunction (10; 11), but the role that vascular changes play in the progression of the disease is widely debated (12; 13). Likewise, a hallmark of tumours is chaotic blood vessel growth (14), leading to leaky microvessels that severely hinder drug delivery, whilst promoting cancerous cell migration (15).

While reconstructed networks are critical tools to identify structural changes during pathogenesis, they are limited in inferring the consequences on functional behaviours. Mathematical and computational frameworks can add valuable insights by using digitally reconstructed networks as the structural basis for simulating physiological processes, e.g. blood flow, molecule exchange (6; 16; 17; 18; 19). Such predictions can then be validated against *in vivo* functional imaging, effectively recreating the structure-function relationship of microvascular networks in health and disease (Figure 1A).

**Figure 1:**
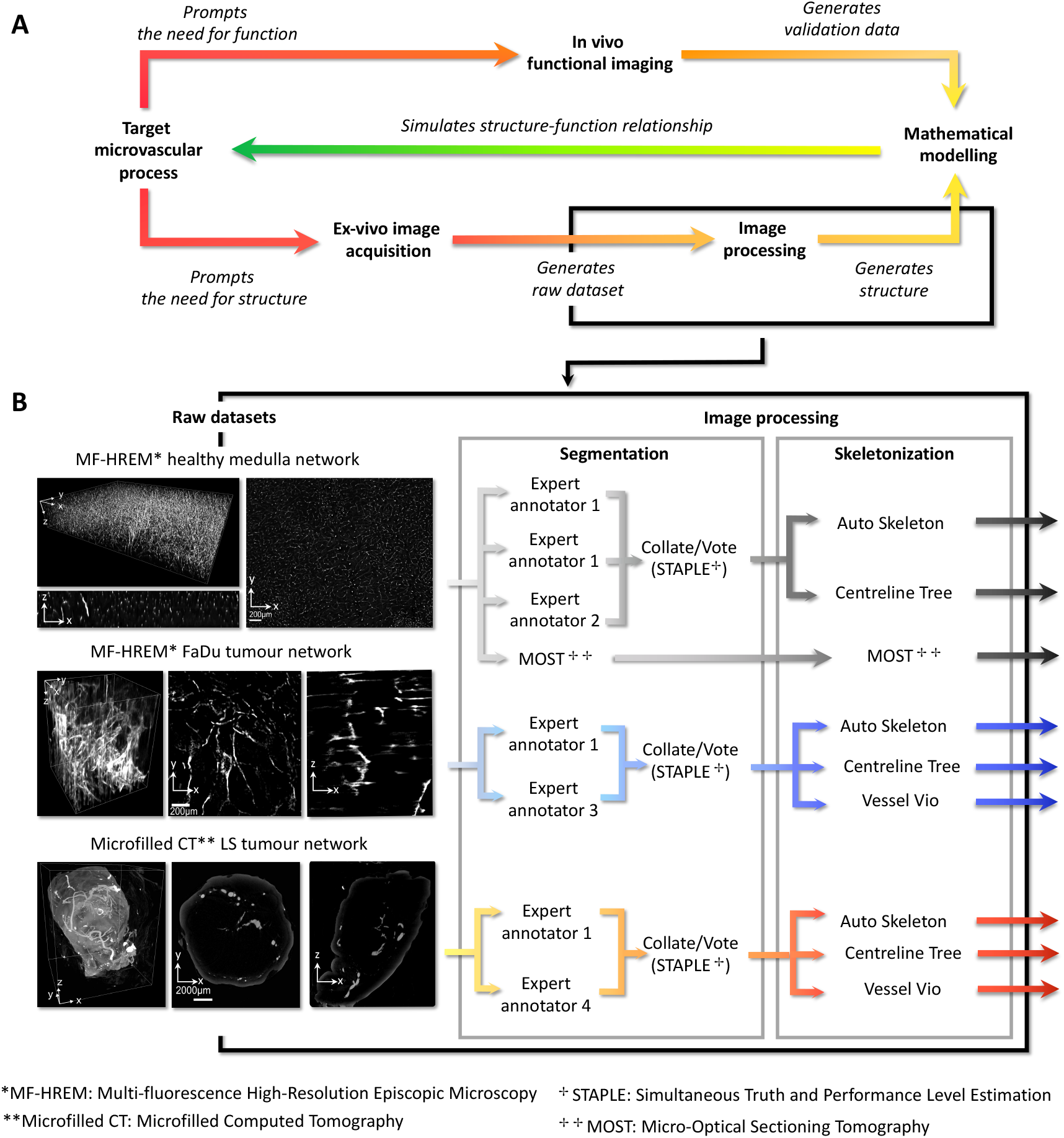
(A) diagram showing the combination of imaging with mathematical modeling to simulate the structure-function relationship of microvascular networks. (B) The image processing section of the imaging-modelling framework, showing the three raw datasets used in this work - Brain medulla, FaDu tumour, LS tumour. Brain medulla and FaDu tumour were acquired using MF-HREM; LS Tumour was acquired using Microfilled CT. Segmentations by individual expert annotations collated via STAPLE algorithm and MOST pipeline (see Section 3.1.2). Skeletonisation of STAPLE segmentations done with Auto Skeleton, Centreline Tree, and Vessel Vio (see Section 3.1.3).

Combining imaging and modelling therefore results in powerful frameworks with the potential to interrogate processes underlying parthenogenesis (16) as well as to accelerate the development of new treatment strategies, through e.g. digital twins(20; 21). However, such frameworks are by nature, composite and therefore are subject to multiple sources of error that can accumulate and propagate from the imaging component to the modelling component, potentially leading to erroneous predictions.

For instance, an inaccurate estimation of vessel diameters during image processing will result in significant variation of blood flow rate predictions, since the microvessel blood flow rates depend on the fourth power of the vessel diameter (22). Blood flow being in turn one of the principal mechanisms underlying oxygen, nutrient and drug delivery, it is therefore critical to be able to estimate the uncertainties associated with the outcome of the imaging section of the framework in order to make quantitative and informative predictions.

The most straightforward approach to quantify such uncertainties is to validate the outcome of each component against a ground-truth (23; 24). Whilst this can be done with physical or digital phantoms, the ability of phantoms to replicate real structures, particularly complex microvascular networks, is limited by both physical phantom manufacture and by the models used to create the synthetic networks (5; 23). In addition, whilst phantoms can provide overall error bounds for a pipeline, they cannot do so for a specific real-world dataset. An alternative approach which acknowledges the lack of the ‘ground-truth’ for a real microvascular network (25; 26; 27) involves creating a gold-standard via a consensus of experts. Such consensus approaches are a widely-applied method in segmentation (28). For spatial graph objects which are the output of skeletonisation, such gold-standard consensus methods are not utilised, making it a vulnerability in the imaging-modelling pipeline.

One reason for this lack of consensus approach in skeletonisation is the challenge of creating suitable metrics for evaluating microvascular networks when represented as graph objects. Graphs contain the spatial location of nodes and segment radii as well as connectivity information for the microvascular network. Averaging or combining this information across multiple spatial graphs, to create an analogue in skeletonisation to the consensus segmentation approach, requires a suitable distance metric to measure between graphs. Whilst some methods have been developed, e.g. using graph edit distances and then looking for a Maximum Common Subgraph (29; 30; 31; 32), the graph edit distances do not provide any way to assess the skeleton’s correctness given the raw image data, or a consensus segmentation. Some metrics for comparison of the raw image data (or segmentation) and skeleton have been proposed, including bifurcation position and number, connectivity, homology etc. (24; 33; 23; 34; 35), however there is no clarity as to which of these metrics is most important or how differences in these metrics translate into differences in the networks functional properties.

Therefore, it is critical to explore and acknowledge how the errors introduced into the image processing pipeline (5; 29; 19; 23) translate into variation in the structural and functional properties of microvascular networks. Here we focus on the variability in microvascular network properties and blood flow predictions that result from implementation of different segmentation and skeletonisation approaches, and delineate a practically implementable super-metric to assess skeleton ‘correctness’.

The challenges to do this are two-fold: 1) to understand the extent to which skeletonisation variation can lead to variation in structural and dynamic network properties; 2) to define a practically implementable metric for validation of skeletons which takes into account salient geometric features. This work addresses these challenges by: 1. Demonstrating the magnitude of uncertainties caused by skeletonisation to both predicted blood flow distributions and structural network measures, 2) Proposing a novel super-metric for skeletonisation evaluation based on comparison of the skeleton to a gold standard binary image used to produce it. Furthermore we develop and introduce this super-metric in a mathematical form that is simple to compute, captures multiple topological characteristics of vascular networks which lead to dynamic functional behaviour discrepancies, and can be formally optimised.

To do this, we utilise three different imaging datasets of vessel networks that bridge different imaging modalities, different scales, and both healthy and pathological networks (see Figure 1B). For each dataset, we create several valid image processing pipelines; that is, pipelines which follow standard methodologies and implement parameterisation methods where these exist. We then compare the outputs of these pipelines in terms of standard structural network metrics, and implement blood flow simulations to gain insight into the link between image analysis tools and flow predictions. This experimental design is summarised in Figure 1B. Imaging meta-data are provided in Supplementary Table S1.

## 2. Sources of variability and mitigation strategies in image processing

Extracting microvascular networks from an image is a two-step process, i) segmentation (Section 2.1), ii) skeletonisation (or meshing) (Section 2.2). We discuss each of these in turn, followed by network structural metrics and the implementation of blood flow models.

### 2.1. Image Processing: Segmentation

Segmentation is the process of partitioning an image into separate segments, e.g. vessel and background, often by assigning a class to every pixel in the image. Segmentation of microvascular networks is a vast research field. Different approaches to microvascular segmentation include filtering for tube-like structures, seed-point growing or flood filling approaches which group connected voxels together, as well as machine learning approaches (36; 37). Validation and benchmarking of segmentation algorithms is done by comparing the output to a ground truth or gold standard segmentation - typically either synthetic data or a portion of manually segmented data (38; 39). Metrics for these comparisons are well-established including overlap based metrics such as DICE or Jaccard indices, surface based metrics such as Hausdorff distances or volumetric based such as Volume similarity etc. (40; 41; 42). More recently metrics which aim to include connectivity as well as voxel overlaps, such as cl-dice (33), have appeared; this metric quantifies what proportion of the voxels that make up the centreline of a segmented vessel fall within the ground truth segmentation. Segmentation algorithms often have many adjustable parameters which are formally or informally optimised, using one or a combination of the above metrics and the ground truth.

Unfortunately, the manual or synthetic gold standard data can also be flawed: synthetic images or images of physical phantoms rarely capture the full complexity of real tissues (28), while manual segmentations, generally performed by expert annotators, are subjective, with factors such as alertness, environmental distractions or differing access to segmentation tools contributing to inter and intra-annotator variation(28; 42). A common approach to mitigate potential discrepancy between gold-standard manual segmentations is to aggregate several manual segmentations through a voting or aggregation strategy. There are several such aggregation strategies of which the most commonly used to for biomedical images is the STAPLE (Simultaneous truth and performance level estimation) algorithm (28; 36; 42; 28; 43; 44).

### 2.2. Image Processing: Skeletonisation

Skeletonisation seeks to reduce the foreground pixels of an image to a thinned (single voxel width) line that largely preserves the extent and connectivity of the original structure. A skeleton can in turn be represented as a spatial graph structure, where meeting points of two or more lines, or end points are nodes and the connections between the meeting points are segments. Such graph structures can serve as the basis for structural analyses as well as simulation of functions such as blood flow.

Different approaches to skeletonisation can be distinguished: 1)thinning approaches which use either morphological operations or distance transformations; 2) minimum cost paths; or 3) wave front propagation. In general the criteria for a correct skeleton given a binary image are: thinness (single voxel thick), medialness (centerline is equidistant from the original boundaries), and homology (a continuous mapping can be made from the 3D volume to the skeleton, i.e. the number of connected components and loops of the original structure is preserved). Thinning approaches are the most common. These iteratively remove pixels of a binary structure without changing the topology (i.e without creating gaps, islands or removal of end points), until a single pixel centreline remains (thinness), which is located at the centre of the original object (mediallness). Medial axis thinning was widely adopted following the approach of Lee et al. (45), which is implemented in many packages (e.g. Matlab, ImageJ, scikit-image and Vessel Vio (46; 47; 48)). Whilst the medial thinning algorithm is highly efficient for 3D volumes it does not preserve homology (49). Alternative thinning approaches were extended by Pudney et al. (50) and Palagyi et. al (51), with application of distance transform ordering (Distance Transformed Homotopic Ordering) (DTHO). These approaches better preserve medialness and have been proven mathematically to preserved homology. DTHO approaches and their derivatives are also widely used and implemented in commercially available packages, e.g. Amira-Avizo Autoskeleton (AS), which implements a parallelised DOHT of Fouard et al. (52).

Minimum cost path approaches such as the Djistrika or Tree spanning algorithms seek to move from one connected node to another - making a (minimal cost) path, through an undirected graph object, i.e. a collection of nodes and edges. These include algorithms such as the TEASAR algorithm (53) implemented in Amira-Avizo’s CentreLine Tree (CL) module.

Wave-front methods simulate flow from initial sets of seed points. The vessel scooping algorithm of Rodriguez et al. (54) adapted by Wu et al. (55) and implemented in an open source framework Vaa3D (56; 57; 58), uses such an approach: starting with seed point(s); at each iteration of the algorithm, voxels within the 3D connected neighbourhood (26 neighbourhood connectivity) of a seed voxel (a single voxel taken as the starting point for the algorithm) are added to make a cluster. At each iteration, a connected component analysis identifies new clusters, which represent a branched vessel, and the centre of mass of every cluster is calculated to define the centerline of the vessels.

These four different methods have been widely applied across different vascular or airway networks(59; 60; 48; 6; 61; 55; 34; 62). Each approach offers different advantages - Computational efficiency is often provided by minimum cost path approaches(53), or parallelised thinning approached (52); better corner preservation is given by the use of distance transformations (50), alongside better delineation of complex junctions etc. (27). Critically, each of these algorithms also applies different constraints on the final topology of the network which will be explored in further detail.

Metrics for comparison and evaluation of skeletonisation algorithms are less widely used or formalised than those used for evaluating segmentation. The most common method found in the literature is to use visual inspection of the skeleton superimposed on the image (52; 45; 63; 34; 27). Some methods or criteria do exist, such as comparison using DICE between manual selection of bifurcation points (64; 65); however, most of these methods focus either on connectivity preservation (a feature that should be assured if homology is preserved) or on volume similarity between the graph and the 3D binary volume. To the best of our knowledge there is no method to rate the correctness of a skeletonisation for a particular dataset which considers both structural and functional properties of the networks.

## 3. Methods

### 3.1. Creation of skeletons

#### 3.1.1. Dataset

We explore three raw datasets acquired using tow modalitied: MF-HREM with fluorescently labeled vasculature, or microCT with microfilled vasculature (i.e. where the vasculature is injected with a radio opaque contrast agent prior to microCT). For each dataset, we applied a consensus segmentation approach using multiple expert annotators followed by skeletonisation using either Amira Autoskeleton (AS), Amira Centreline Tree (CL) or VesselVio (VV), or the MOST algorithm, which is an end-to-end segmentation and skeletonisation method. The nine outputs from these pipelines were compared via structural metrics and a blood flow simulation metric. Figure 1B summarised the pipeline. The three datasets used were:

- Portion of a brain medulla network from a W/T Balb/c mouse, collected with MF-HREM (8))
- Portion of a subcutaneous FaDu tumour network from a Balb C mouse imaged with MF-HREM (8)
- Whole subcutaneous colorectal tumour (LS174T) (referred to hereafter as LS) imaged with microCT after microfill filling on the vessels (66).

Raw datasets are available via the respective publications.

#### 3.1.2. Segmentation

Manual segmentation by two or more annotators (referred to as ‘Expert annotator’ in Figure 1B) were performed on all raw datasets using AmiraAvizo v2019.1-2021.2. The segmentations were performed using a manual local region filling tool (magic wand). Using this tool, an annotator selects seed point(s) in any one of 3 orthogonal image planes, as well as selecting and varying intensity and contrast thresholds; voxels connected in 3D to the seed point which are within an annotator set threshold for intensity or contrast, can then be selected and added to the segmentation. In addition a physical boundary or limit could be drawn by the annotator to limit the extent of the region growing, and manually corrected by voxel painting where can be used to correct the segmentation on individual slices. In this way it is possible to for a trained annotator to manually segment relatively large 3D vascular datasets without being biased by the plane of the image slice they are segmenting in (as created by interpolation methods in many manual approaches)

A total of four annotators participated; all, barring annotator 2, could be considered as experts for the specific imaging modality. The STAPLE algorithm, an iterative weighted voting algorithm (28) was applied to the expert annotations for each dataset. At each iteration this algorithm votes on every pixel in an image segmentation based on the set of annotations provided by the experts. It then weights each annotator according to how closely their segmentation corresponds to the voted image. This weighting is then used in the next iteration of the voting process. The algorithm continues until the segmentation stops changing. This process creates a single segmentation for each dataset that is considered to be a consensus segmentation between the expert annotators.

Widely used metrics were applied to compare each individual expert manual segmentation to the consensus segmentation. Metrics recorded were Jaccard index, Dice score, Volume Difference and Haunsorff Distance:

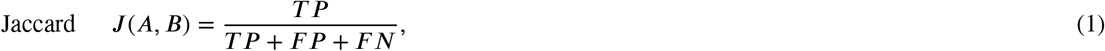

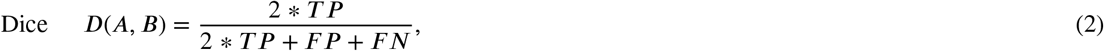

where *T P* is the voxel-wise true positive (the number of voxels where a vessel is detected in both the consensus and test segmentation images at the same location), *F P* is voxel-wise false positive (the number of voxels identified as a vessel in the the test segmentation but not in the consensus) and *F N* is the voxel-wise false negative (the number of voxels where a vessel is not detected in the test segmentation but is in the consensus segmentation). Unused in these metrics is the voxel-wise true negative, *T N*: the number of voxels where a vessel is not detected in both the consensus and the test segmentations.

Volume difference is a typical volume-based metric that is based on the absolute difference in volumes of the segmented structure and the ground truth, usually normalised by the ground truth volume:

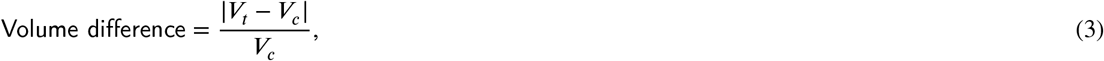

where *V*_*t*_ is the volume of the test segmentation and *V*_*c*_ is the volume of the consensus segmentation.

The Hausdorff distance is commonly used to compare two voxelised surface representations e.g. to compare boundary points from the consensus segmentation *A* and test segmentation boundary points *μ*. It is defined as the maximum distance between each point in *A* to its nearest neighbour in *μ*.

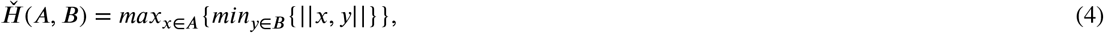

Where 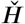 is the directional Hausdorff distance from which the absolute Hausdorff distance *H* can be calculated.

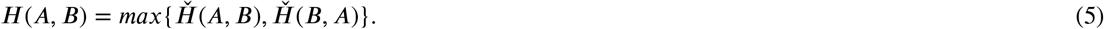

#### 3.1.3. Skeletonisation

Skeletonisation reduces the network to a graph representation which describes the vessel network in terms of ‘nodes’, ‘points’, ‘segments’, and ‘sub-segments’ (illustration for each of these structures are provided in Supplementary Figure S1). A segment is defined by a start (*i*) and end node (*j*) (these nodes have an ID and a 3D spatial position (x,y,z)); which could either be a branching node connecting several segments together or a terminal node where no further branches were detectable. Between the start and end node of each segment lie sub-segments (*s*), with ‘points’ marking the start and end of each sub-segment, able to capture the curvature of the segment. Each sub-segment has an associated radius (*R*_*s*_) and length (*L*_*s*_)).

We implement and compare a number of skeletonisation algorithms chosen according to the following criteria: i) we chose not to apply any in-house codes that do not have a well maintained open-source platforms, ii) the algorithms should be able to run on large (<100GB) image volumes, iii) they should have been used by other groups to produce segmentations of large blood vessel networks, iv) our choices should cover the range of methodological approaches outlined in Section 2.2.

Using these criteria four skeletonisation methods were chosen, three of which were run with binary image inputs and one which used raw image data as an input:

- Amira v2021.2 Autoskeleton (AS) plugin: a widely used skeletonisation package (6; 62; 67; 34), which implements the a parallelised version of DOHT algorithm (52). Amira-Avizo’s implementation estimates the radius of each subsegment using 1/5th of the maximum Chamfer distance and provides additional optional user inputs to smooth the output of the spatial graph. If smoothing is selected, there are two associated parameters, ‘smooth’ and ‘attach to data’, smoothing is done via weighted average of a point’s location and the location of its two neighbours; the ‘smooth’ and ‘attach to data’ provide the respective weights this averaging. This smoothing can be performed iteratively and the iterations can also be defined by the user. Finally, a threshold value is available for creating a binary image if the user input has not already been segmented (for a binary image this is automatically detected).
- Amira v2021.2 Centerline Tree (CL) plugin: an implementation of the TESEAR algorithm (53), a shortest distance path spanning approach which creates a strict tree structure. There are two user-defined parameters: the zeroVal and slope parameters, which are used to restrict the voxels which are included in the search for end points of the network, For any particular segments centerline, any voxels which fall within the critera: slope ^*^ distance to boundary of vessel + zeroVal are excluded from the end point search. This reduces the sensitivity of the algorithm to surface noise in the segmentation. The radius is estimated using a minimum inscribed sphere algorithm(53). CL has been used to skeletonise various vascular networks (60; 59; 68; 69).
- VesselVio (VV): a recently released open-source package (48) for skeletonisation of large vessel networks. It provides an implementation of the widely used medial thinning approach of Lee et al. (45), with automated calculation of common structural metrics. In this case modifications have been developed to detect spurious branches and to allow pruning of structures. These are controlled by three user inputs: length of end segments to be pruned, length of isolated segments to be pruned and resolution of the image. The subsegment radii are estimated using a Modified Euclidean Distance map (48).
- Micro-Optical Sectioning Tomography (MOST): an end-to-end pipeline, i.e., it does not require a binary input image and instead can take a raw image data as an input. It performs a vessel scooping algorithm (54) adapted by Wu et al. (55). Starting with seed point(s); at each iteration of the algorithm, voxels that are above a threshold and are within the 26 neighbourhood connectivity of a seed point are added to make a cluster. At each subsequent iteration, any voxels above the threshold, that are connected to the cluster and within the scooping radius (parameter set by user), are added to the respective cluster. A connected component analysis on the clusters at each iteration identifies new clusters (i.e. those not connected to each other) which delineate a branched vessel. The centre of mass of every cluster is calculated at each iteration to define the centerline of the vessels. The radius of the vessels is calculated using a radius-adjustable sphere algorithm, where radius of the vessel is equal to the radius of the maximum sized sphere that fits entirely within the segmented vessel (58). This has been implemented in the open-source Vaa3D software (56; 57; 58) and requires 4 user defined parameters: the threshold, the seed size, the scooping distance and the size of the voxels (non-isotropic voxels are supported). This pipeline has been applied to mouse brain vessel segmentation for the MOST imaging technique (55). Here, this method was applied to only the medulla network.

The selection of parameter values for each skeletonisation algorithm is challenging when applying to large complex network structures due to a lack of defined metrics which will be discussed later. For AS, CL and VV algorithms, parameters were set via overlaying the skeleton on the image volume and visually inspecting the data for poor overlap, including addition of spurious nodes or segments, or missing portions of network (See Supplementary Table S1 for values). For the MOST pipeline, an existing methodology for parameter setting developed by (55) was followed and is outlined in detail in the Supplementary information section 1.1.

### 3.2. Analysis of skeletons

#### 3.2.1. Structural metrics

Many different structural metrics are used to characterise and compare microvascular networks. Table 1 lists a number which are used extensively in the literature, organised into sub categories based on similar properties. These categories are: vessel diameter and volume measures, branching measures, vessel spacing and density measures, and network-shape measures.

**Table 1.** Morphometric measures of blood vessel networks presented in the literature. These are metrics that depend on vessel diameter and volume, vessel length measure, branching parameters and vessel spacing and density measures.

| Structural Metric (as named by authors) | Description | References |
| --- | --- | --- |
| **Vessel diameter and volume metrics** - Vessel diameter varies substantially throughout vessel networks and affects blood flow and pressure. It has also been used as an identifier of pathology (70; 71) |  |  |
| Vessel diameter | Vessel diameter | (6; 72; 73; 74; 75) |
| Average radius | Average vessel radius | (34) |
| Number of dilated capillaries | Number of vessels with a diameter above 40μm | (71) |
| Variation from mean vessel diameter | Percentile variation from mean vessel diameter | (70) |
| Length-to-diameter ratio | Length to diameter ratio | (76) |
| Vessel volume | Total volume of all vessels can be calculated from area and lengths of all vessel subsegments | - |
| Compactness | Ratio of area/volume of convex hull containing vessels (polygon enclosing all vessel) to the total area/volume of all vessels (density type metric) | (77) |
| **Branching metrics** - These look to quantify the properties of bifurcations within networks, which have been shown to differ between healthy and pathological networks (72). Branching metrics also have a direct bearing on predictions of flow and oxygen distributions, as hematocrit is unevenly split between daughter vessels based on the flow in each of them. |  |  |
| Branching number | Number of daughter vessels connected to a parent at a branching node | (6; 72; 73) |
| Branching angle | Angle between two vessels at a branching point | (6; 72; 73) |
| Branching order | The level in the hierarchy of a branching structure a vessel is located, with outermost vessel in this case considered as 1st Order. | (75) |
| Bifurcation index | Ratio of diameters of two daughter vessels at a branching point | (78) |
| Branching index /bifurcation density | Number of branching points per unit area/volume | (79; 34) |
| Ratio of length of daughter to parent | Ratio of length of daughter vessel to parent vessel | (78) |
| Interior node density | Number of non-boundary nodes per volume | (74) |
| Boundary nodes | Number of boundary nodes per surface area of the region of interest | (74) |
| % multiply-connected nodes | Percentage of interior nodes with more than 3 connections | (74) |
| **Vessel spacing and density metrics** - All cells in the body require oxygen and the intervessel distance is an indicator of the diffusion distance from blood vessels to cells. Large intervessel distances are indicative of oxygen-deprived regions in the tissue which can often be markers of pathological cell function or can identify different microenvironment niches. |  |  |
| Intervessel distance | Distance between vessels | (6; 72; 73) |
| Minimum intervessel distance | Minimum distance between vessels | (75) |
| Vessel density | Vessel volume/ Whole volume | (6; 72; 73; 80; 79; 81) |
| Interbranch distance | Distance between branching points (i.e. vessel length) | (6) |
| Highest microvascular density | Highest vessel volume/whole volume | (80) |
| Maximum extravascular distance | Maximum distance between a vessel and a tissue point | (74) |
| Convexity index | Slope of linear fit to log-log scale histogram of extravascular distance (inversely correlated with maximum extravascular distance) | (74) |
| **Network shape metrics** - these provide information on structures in the network including the number of loops, the number of edges per vessel loop and the loop length. Computational complexity associated with most of these metrics is high (74), with the exception of tortuosity. Blood vessel tortuosity is a sign of pathology, for example in oncology, ophthalmology, cardiology and neurology (76). Tortuosity may be calculated in many ways including the distance metric, the Sum of Angles Metric (SOAM) or the inflection count metric (ICM). |  |  |
| Tortuosity | The ratio of the curved to straight line length of the vessel | (76) (82) |
| Number of loops | Number of loops created by vessels | (75) |
| Mean no edge/loop | Number of edges per vessel loop | (74) |
| Mean loop length | Mean loop length | (74) |
| Total number of endpoints | Total number of branch endpoints (e.g. boundary or deadend) | (79) |
| Number of Subgraphs | The number of subgraphs within the network | (17) |

As Table 1 indicates there are many metrics to choose from when analysing microvascular networks See Supplementary Figure S1 for additional diagrams. Here we choose a subset that are frequently reported in the literature, will provide comparative insights across our three networks and have a bearing or link to the network functionality:

- Radius – 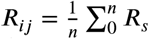 where *n* denotes the number of subsegments comprising the segment {*ij*} linking node *i* with node *j*.
- Length to Diameter Ratio – 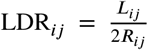 where *L*_*ij*_ 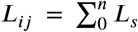 denotes the length of the segment, i.e., the cumulative length of each subsegment.
- Tortuosity – 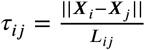 where ||***X***^*i*^ −***X***_*j*_ || represents the Euclidean distance between ***X***^*i*^ and ***X***_*j*_ the positions of node *i* and *j* respectively.
- Number of sub-networks - *cc* the number of distinct sub networks, i.e. graphs not sharing a single node with one another.
- Branching Angle – 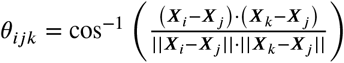 the direct angle between segment *ij* and segment *jk*
- Intervessel Distance – 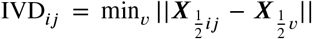 where 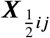 represents the position of the midpoint of segment *ij* and 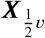 represents the position of the midpoint of any other segment, so that IVD is the closest midpoint to midpoint distance to another segment, for every segment.

These metrics are summarised in Figure 2 and diagrammatically shown in Supplementary Figure S1.

**Figure 2:**
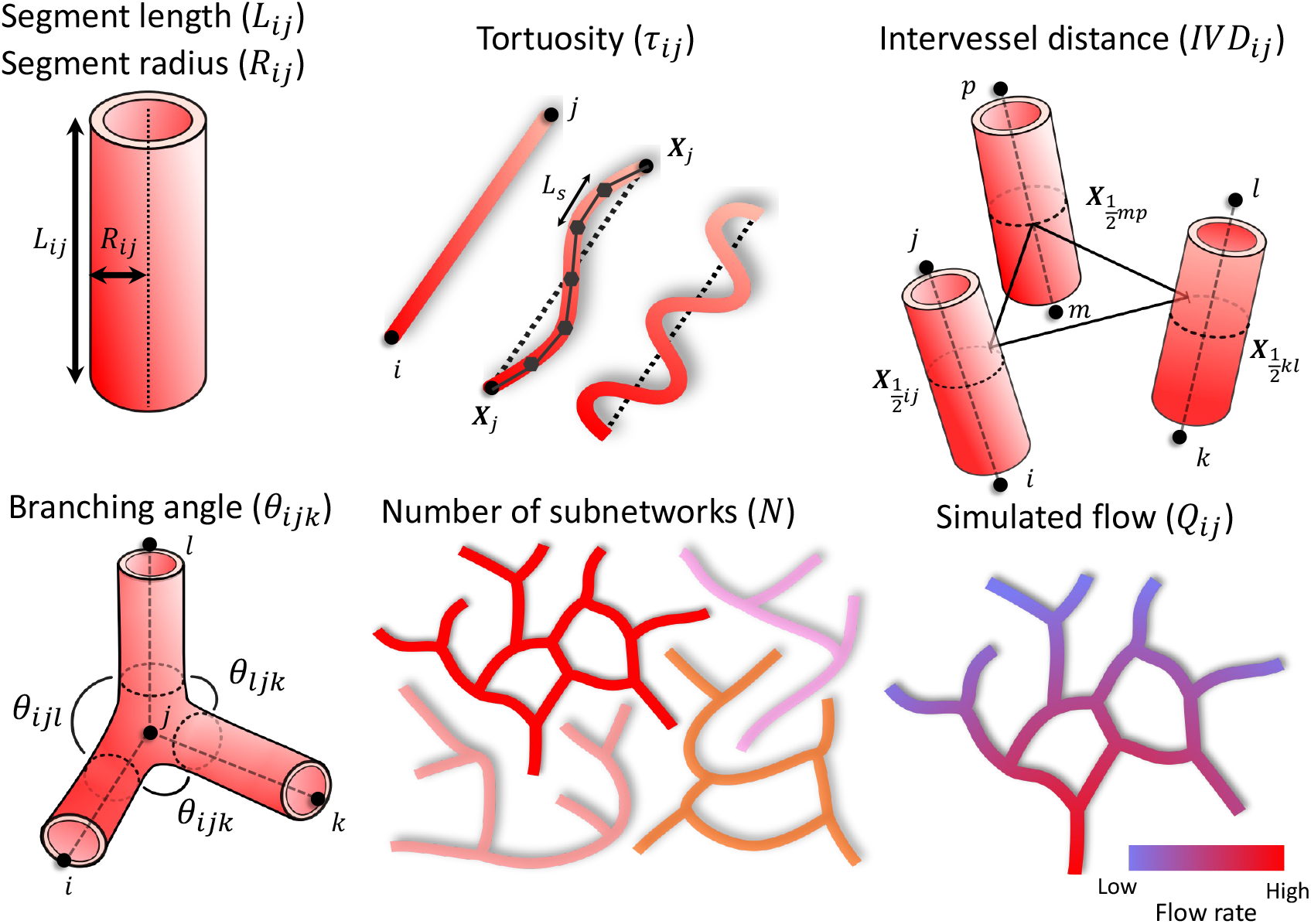
Categories of metrics computed for each network (medulla, FaDu, LS) and for each skeletonisation algorithm (MOST, AS, VV, CL). See Supplementary Figure S1 for diagram.

#### 3.2.2. Blood flow simulations

A range of approaches exist to simulate blood flow in microvascular networks, from fully-resolved three-dimensional implementations suited for small networks (83), to continuous formulations describing flow distribution at the scale of entire organ (84; 85). In this work, we focus on a popular pore-network approach (6; 17; 86; 10) that does not requires further processing and makes direct use of the network skeleton (indeed, this approach has already been used in imaging-modelling pipelines (16)). We describe blood as an effective fluid with non-Newtonian properties and assume Poiseuille flow, reducing the problem to a pressure distribution associated to the nodes of the graph, and flow rate distribution associated to the segments of the graph. First, flow rate conservation is imposed at every node of the graph so that

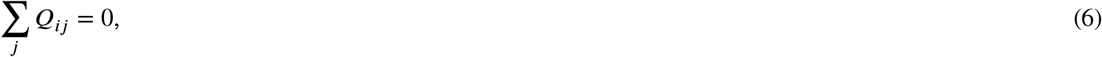

where *Q*_*ij*_ represents the flow rate in segment *ij*, where *i* and *j* are the nodes at each extremity of the segment. Then, momentum conservation is prescribed between those two nodes assuming no leakage

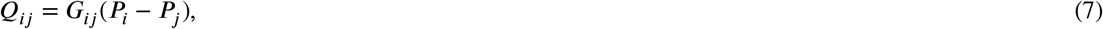

where *P*_*i*_ and *P*_*j*_ represent the pressure at node *i* and *j* respectively. Here *G*_*ij*_ represents the conductance associated with the segment *ij* and is defined as

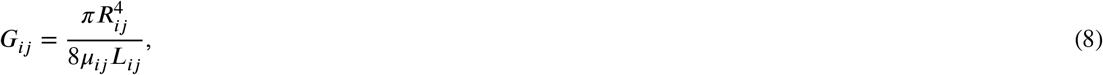

where *R*_*ij*_ is the segment radius, *L*_*ij*_ the segment length accounting for segment tortuosity (*τ*_*ij*_ ) and *μ*_*ij*_ to the blood effective viscosity. Such a viscosity depends non-linearly on both the segment diameter and local presence of red blood cells, referred to as the haematocrit, and is described using a well-established relationship derived from experiments (87). In this work, we consider that each segment has a constant haematocrit corresponding to the systemic haematocrit (i.e. *H*_*d*_ = 0.4 (88)), although this could be readily extended using additional semi-empirical relationships to account for phase separation effects (89), with the trade-off of solving a coupled, non-linear problem.

We close the system formed by Equations (6)-(8) by applying boundary conditions at all terminal nodes, i.e. nodes connected to exactly one segment. The assignment of boundary conditions in the absence of measured data is highly challenging - here we seek a pragmatic approach which allows us to explore the role of vessel architecture on network-scale functional metrics (rather than being able to make specific, quantitative predictions). Therefore we develop an approach which is straightforward to implement and standardisable across the different networks:

- The Medulla and FaDu networks have a box-like shape, with a large number of terminal nodes near the box faces. We apply a pressure drop Δ*P* = 50mmHg between opposite faces, observing that the blood flow model (Equations (6)-(8)) is linear so that blood flow distribution is independent of pressure drop. We chose this value so as to remain in a physiological range (10) and as a basis for discussions. We assume terminal nodes located within 50 microns of a face belong to that face and that terminal nodes located close to the box corners were associated with their closest face. For nodes lying on other faces we impose 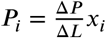 where Δ*L* is the distance between the two opposite faces and *x*_*i*_ the position of the node along the direction of the pressure drop. For terminal nodes located within the network, i.e. dangling ends located far from the faces, we impose no flow, so that inlet and outlets are located only on the network surfaces. To avoid favouring a specific direction we repeat the simulations with the pressure drop applied on the remaining two pairs of opposite faces. See Supplementary Figure 3 for diagrammatic representation.
- The LS network includes structures with larger diameters than the other two networks and is also less box-like in shape. In particular, Figure 5vii shows that the LS network has a large segment dangling close to the bottom. This segment is present in every reconstructed skeleton, regardless of the algorithm considered. The terminal node associated with such a segment is then prescribed with a high pressure, and all other boundary vertices are prescribed with a low pressure so that the pressure drop between high and low pressure vertices is Δ*P* = 50mmHg.

**Figure 3:**
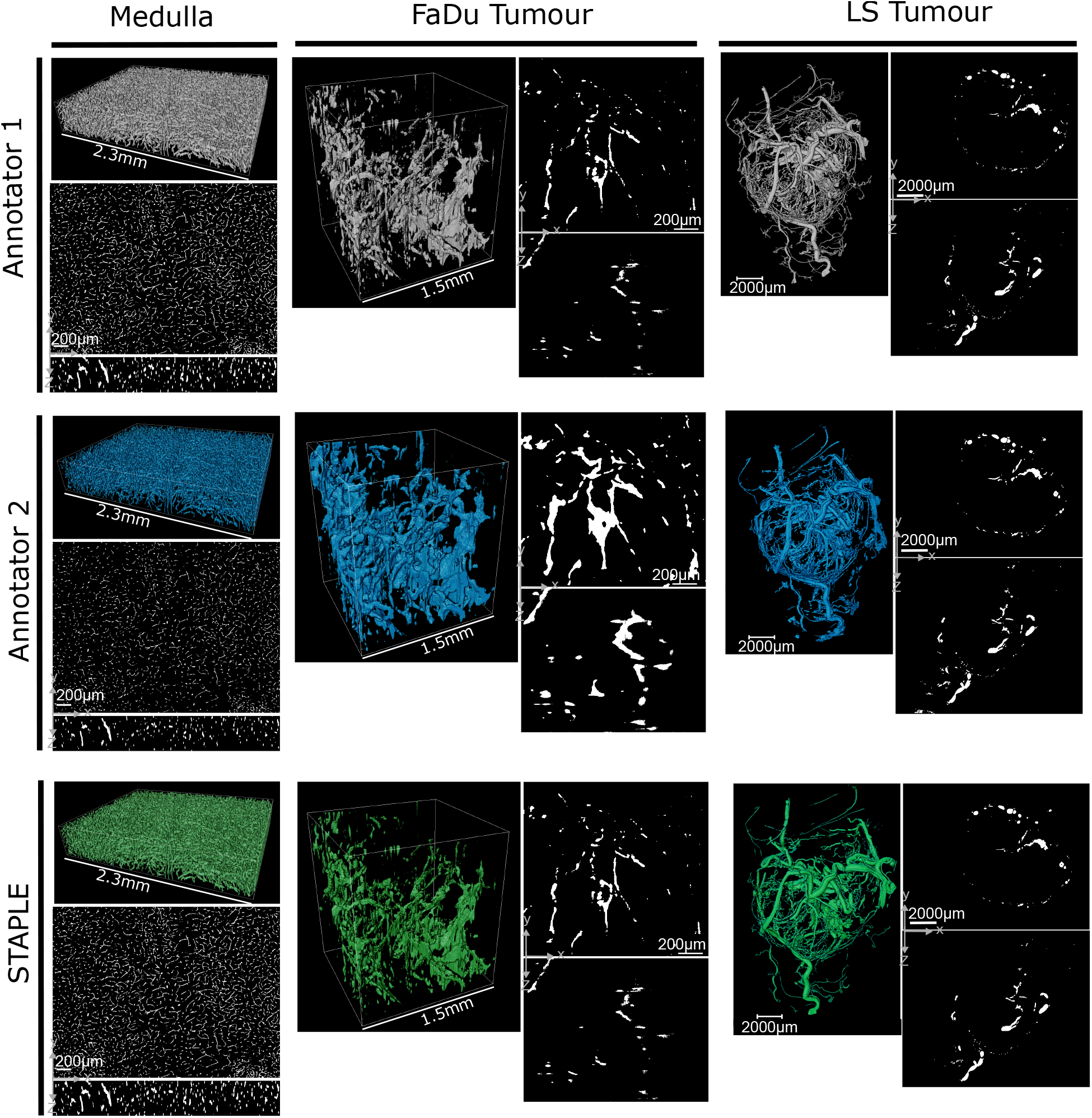
Segmentation outputs for each dataset from two independent annotators. The bottom row shows the STAPLE output for each dataset.

We perform blood flow simulations for the Medulla, FaDu and LS networks for each image processing pipeline presented in Figure 1B.

## 4. Results and Discussion

### 4.1. Segmentation

The segmentation metrics for the manual annotator segmentations of each dataset, which compare the overlap between each expert annotator and the consensus STAPLE segmentation, are provided in Table 2. In all cases, the different annotators produced substantively different segmentations from one another, with one segmentation being quantitatively similar to the STAPLE consensus.

**Table 2.** The Jaccard index, Dice Score, Volume similarity and Hausdorff Distance, of individual semi-manual segmentations compared to a gold standard segmentation. The gold standard was constructed by combining the individual hybrid hand-automated segmentations using the STAPLE algorithm.

| DataSet | Annotator | Jaccard Index | Dice Score | Volume Similarity | Hausdorff Distance |
| --- | --- | --- | --- | --- | --- |
| MR-HREM Medulla Brain | Non-Expert Annotator | 0.52 | 0.68 | 0.6322 | 82.32 |
| MR-HREM Medulla Brain | Expert Annotator (1) | 0.66 | 0.80 | -0.40757 | 55.0 |
| MR-HREM Medulla Brain | Expert Annotator (1) | 0.99 | 0.99 | 0.000137 | 57.2 |
| LS Tumour CT | Expert Annotator (3) | 0.84 | 0.91 | -0.171 | 83.21 |
| LS Tumour CT | Expert Annotator (1) | 0.74 | 0.85 | -0.31 | 72.01 |
| FaDu Tumour MF-HREM | Expert Annotator (4) | 0.28 | 0.43 | -1.13 | 100.3 |
| FaDu Tumour MF-HREM | Expert Annotator (1) | 0.99 | 0.99 | -0.006 | 77.6 |

The Medulla brain network represents the simplest segmentation case - vessels are well labelled throughout and imaging artefacts (anisotropic asymmetric resolution specific to the imaging modality MR-HREM) have been well reduced (90) Figure 3. Additionally, the network is non-pathological and thus vasculature has a more ordered structure (i.e. vessels appear tube-shaped with most branches have one parent and two daughter nodes).

The FaDu and the LS tumour networks both represent pathological microvascular networks, with highly disorganised vasculature where many vessels do not appear tube-like in structure, and complex branching points where many vessels meet Figure 3.

The LS tumour annotators performed their segmentations Figure 3 in approximately 120hrs (Annotator 1) and 15hrs respectively (Annotator 3). Annotator 1’s segmentation shows better distinction of the fine vessels and this is reflected in the metrics in Table 2. This highlights that not only is the level of expertise important to consider, but also the conditions under which each expert annotator is working. As of yet, there is no widespread consensus for the best approach to these challenges (91; 42).

The FaDU network represents the worst case scenario for segmentation: the network is pathological, meaning deviations from tubular branching tree structures cannot be used to distinguish imaging artefacts from vascular structures. Also the anisotropic resolution, characteristic of MF-HREM (90), is more pronounced owing to the lower resolution Figure 3. These challenges are borne out in the differences between the two expert annotators (Table 2). In this case experts had had similar levels of experience and completed the task over similar time frames.

The MOST algorithm combines both segmentation and skeletonisation, and utilises a different form of adaptive threshold for the segmentation portion of the algorithm. The segmentation portion of this algorithm was not evaluated separately to the skeletonisation portion, rather the final output was evaluated against the STAPLE segmentation using overlap metrics, precision and recall, defined by the authors of the MOST algorithm (55). We combine the precision and recall by taking the harmonic mean of these and refer to this as the F1 score (see Supplementary Materials Section 1.1). Our highest score was F1=0.83. This cannot be directly compared to the overlap metrics of DICE or Jaccard indices of the manual segmentation in the previous sections, as it is based on the overlapping ‘regions’ of vessel in cross-sectional images throughout the stack, rather than voxel-to-voxel overlap on the full image volumes, and only after a re-binarization of a skeletonised graph. Rather, the F1 score we obtain should be compared to 0.92, which is calculated from the harmonic mean of precision and recall in (55). The lower F1 score here already suggests that there will be high uncertainty in the final skeleton results for this algorithm.

In all cases it is clear that whilst there may be closer or further consensus between experts regarding the manual segmentation of microvascular structures, one should be cautions of inferring ground-truth from consensus, and should rather consider gold-standard consensus as a distinct case from e.g. a synthetic dataset ground truth.

### 4.2. Skeletonisation metrics

For all datasets each skeletonisation results in a spatial graph with the nodes, segments etc. as previously discussed. The number of nodes, segments, subnetworks, total network volume, branched and terminal nodes are reported in Supplementary Table S3. From these data alone, the wide variation produced by these different methods is clear. For each dataset, the skeletonised network with highest number of nodes has more than double the number of nodes and segments than that with the lowest. In addition, there is no clear pattern for any given algorithm: e.g. for the tumour datasets, LS and FaDu, the CL algorithm produces the lowest number of branching nodes, whereas for the medulla brain dataset the CL algorithm produces the highest number of branching nodes. This could be attributed to the number of looping nodes that are broken by the CL algorithm which the medulla may have more of given that it is the highest resolution dataset. Comparison of the network volume with the volume of the consensus segmentation (Supplementary Table S3-S5) demonstrates VV provides the closest volume approximate in the FaDu and LS cases, which can be linked to the mean radius result in the structural metrics we now discuss.

The structural metrics comparison demonstrates the differences in the geometry of the networks using metrics which are commonly applied in the field (section 4.2). Figures 4 and 5 show the distributions of structural metrics for the spatial graphs produced by skeletonisation of the STAPLE segmentation and MOST output for all three datasets. The radii, LDR, branching angle, IVD, number of subnetworks and tortuosity for each of the skeletonisation approaches are statistically compared based on the differences in respective distributions (as measured by the Kruskall-Wallis test) and are summarised in Table 3 (extended statistics in Supplementary Table S10-S12).

**Table 3.** Summary statistics for Kruskall-Wallis test between means of each structural metric across each skeletonisation method, *** p *** 0.0001, ** p ≤ 0.001, * p ≤ 0.05, ns p ≥ 0.05

| Datasets | Radius | Branching | IVD | Tortuosity | LDR | Flow |
| --- | --- | --- | --- | --- | --- | --- |
| **Medulla** |  |  |  |  |  |  |
| MOST vs. AS | *** | *** | *** | *** | *** | *** |
| MOST vs. CL | *** | ** | *** | *** | *** | *** |
| AS vs. CL | *** | *** | *** | *** | *** | *** |
| **FaDu** |  |  |  |  |  |  |
| VV vs. AS | *** | ns | *** | ** | *** | *** |
| VV vs. CL | *** | ** | *** | *** | ** | *** |
| AS vs. CL | *** | ** | *** | *** | *** | *** |
| **LS** |  |  |  |  |  |  |
| VV vs. AS | *** | ns | *** | ** | *** | *** |
| VV vs. CL | *** | ns | *** | *** | *** | ns |
| AS vs. CL | ns | ns | *** | *** | *** | *** |

**Figure 4:**
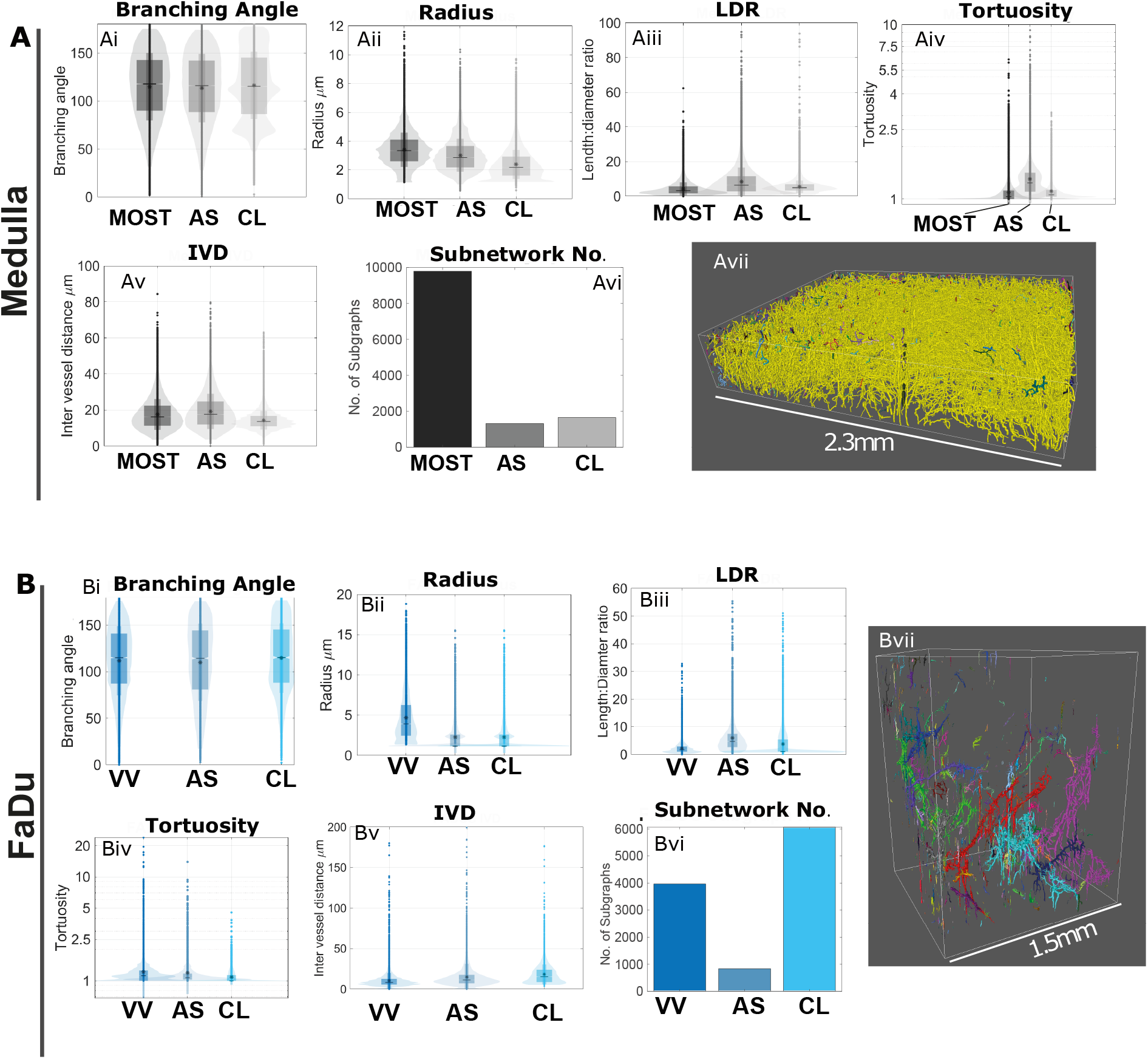
(A) Medulla brain and (B) FaDu tumour network metrics. For both cases i-vi are the branching angle, radius, LDR, tortuosity, IVD and number of subnetworks respectively, and vii shows the network with subnetworks coloured for one of the three image processing pipelines.

By looking across all datasets, the algorithm dependency of structural metrics becomes evident in some cases. For example, in all cases the skeletonisation algorithm that produces the highest mean radius value produces the lowest IVD. This is VV in all cases where it is applied and can be seen to be a consequence of the modified Euclidean distance approach to radius estimation (48). Similarly the networks with the highest number of subnetworks tends to have the highest IVD, although this can be countered by large radii. The majority of the metrics differ significantly between skeletonisation methods. Considering the structural metric results in terms of the skeletonisation methods: the CL algorithm, a shortest path approach following a distance ordered search, produces a less well connected network (higher number of sub networks) compared to the thinning approaches of VV and AS (Figures 4Bvi and 5vi). The AS algorithm consistently produces the lowest number of sub networks across all datasets, though it is not clear if this is a ‘better’ skeletonisation, we discuss this notion further section 5. For other metrics, there is no clear pattern in the variation between CL and AS approaches across all datasets. For example, the mean IVD in the Medulla brain network is significantly lower in the CL case than in the AS case, whereas the reverse is true for the FaDu network. For tortuosity there is also not a clear pattern in the differences between the distributions across all three datasets (Figures 4Aiv, 4Biv and 5iv).

**Figure 5:**
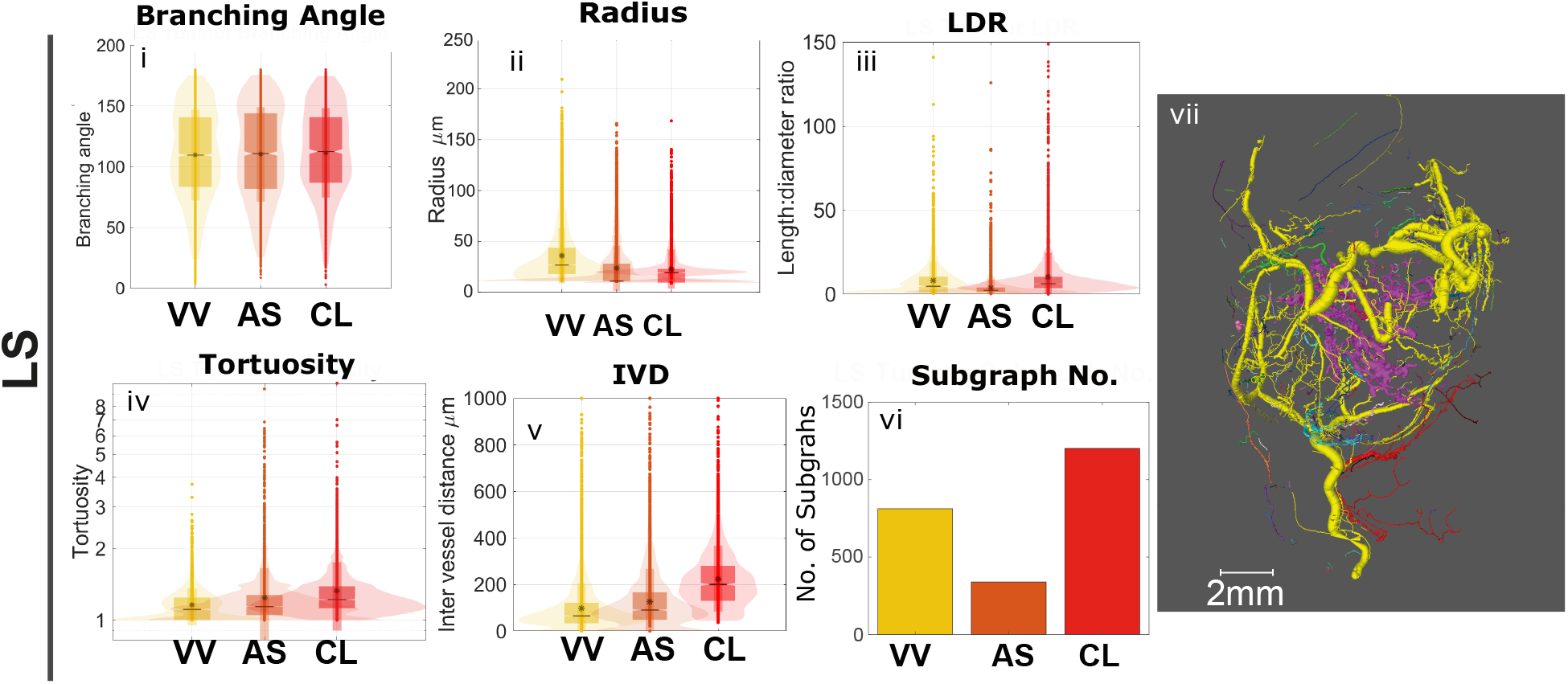
LS Tumour network metrics. Graphs i-vi are the branching angle, radius, LDR, tortuosity, IVD and number of subnetworks respectively, and vii shows the networks with subnetworks coloured for one of the three image processing pipelines.

Other metrics, such as the radius distribution, display some patterns that can be linked back to assumptions or limitations of the algorithms; for example, we note that the radius distribution for the CL has a hard-coded minimum radius value of twice the pixel size. This distorts the radius distribution where vessels approach the imaging system resolution (Figures 4Ai, 4Bi and 5ii). Where the imaging resolution is lower, and thus the image captures mostly larger vessels (such as the LS tumour dataset), the hard coded minimum is less apparent (Figure 5i).Similarly the radius calculation for the AS algorithms has a simplistic approach which uses 1/5th the maximum of the Chamfer distance map. This lead to generally lower radius values than other algorithms for the lower resolution datasets which use e.g. maximum inscribed sphere approaches used in the CL (53) and MOST (58) or a modified Euclidean distance approach in VV (48). In addition, the strict tree topology enforced by the CL algorithm, and the MOST breaks looping structures within the network forcing the structure towards a more ‘tree-like’ topology i.e single parent vessel with two child vessels at nodes. This leads to a higher number of terminal nodes compared to all other skeletonisations algorithms (Supplementary Table S3). In the CL case this also leads to distinctive bimodal distribution of branching angles with the smaller angle between child vessels and a larger the angle between parent and child. This forcing of networks into a strict tree-like topology may be appropriate for large calibre vessels or in networks that are purely an arterial or venous, but is a highly questionable assumption in the case of capillary networks which have a more mesh-like structure (17) such as the medulla brain. Moreover, is is not clear that such an assumption would hold for a pathological vessel networks such as the tumour networks of the LS and FaDu. The MOST approach, the only approach which does not share the same binary segmentation, produces even further disparity in metric results, including a reduction in network connectedness, as evidenced by the high number of subnetworks (Figure 4Avi). This is likely influenced by the absence of manual segmentation in this pipline; human annotators have a tendency to fill in connected regions when segmenting even if there is faint or absent staining, due to the expectation of a connected tubular structure when segmenting a microvascular network (36). The differences between network topologies for the AS and VV approaches are of particular interest (Figure 4B and Figure 5) as both are thinning approaches with the same input segmentation. This shows that these widely used algorithms with the same theoretical aims (e.g. medialness, homology, and thinness) result in significantly different graph outputs. Furthermore it is not possible to visually assess the output correctness through overlay methods alone, or to use the quantitative structural metrics of the outputted networks to determine which algorithm performs ‘better’ in either dataset. Indeed, neither of these methods produces a homotopic skeletons which is shown simply by comparison of the number of connected components in the binary labelled volume and the number of subnetworks (see Supplementary Tables S3 - S5 for values). Interestingly, the CL algorithm does preserve the number of connected components when compared to the input segmentation. However, as previously discussed, it does not preserve loops and thus will inevitable violate homology where looping structures are present in the segmented micro-structure.

In some cases the comparison of the structural metrics does not achieves statistical significance, e.g. in the LS network branching angle (Figure 5i). This could suggest that two algorithms are in closer agreement than a third which differs significantly; and whilst this may be the case, a spatial encoding of the branching angles (as with the other metrics) is necessary to understand if structural differences will result in a functional difference between two skeleton structures. This is something which is rarely considered in spatial graph microvascular network analysis, but can be born out through functional metrics such as the flow results we present below.

Overall the structural metric analysis highlights significant variability underpinned by skeletonisation algorithm selection. Whilst we do not question that structural metrics do vary between networks in meaningful ways such as in pathology and as such, are still clearly valuable to compute. We show that great care must be taken to ensure that the measures extracted are not dominated by the skeletonisation algorithm used to compute and moreover that any comparison of networks using different skeletonisation algorithms such as between different studies, should be treated with extreme caution. It is also clear that structural metrics alone do not necessarily provide a clear indication of whether any one skeletonisation algorithm is closer to an accurate representation of the imaged microvascular networks than another.

### 4.3. Flow results

The flow model used here incorporates geometrical and topological network information with a standardised approach to boundary condition assignment; therefore, we consider the flow predictions as a summary statistic on how different structural interpretations of the same networks would influence functional measures rather than a quantitative prediction about *in vivo* dynamics. In this context, Figures 6 and 7 show the simulated flows across the different skeletonised networks (VV, AS, CT and MOST), with Figure 6A and B focusing on the medulla and FaDu networks, and Figure 7 on the LS network.

**Figure 6:**
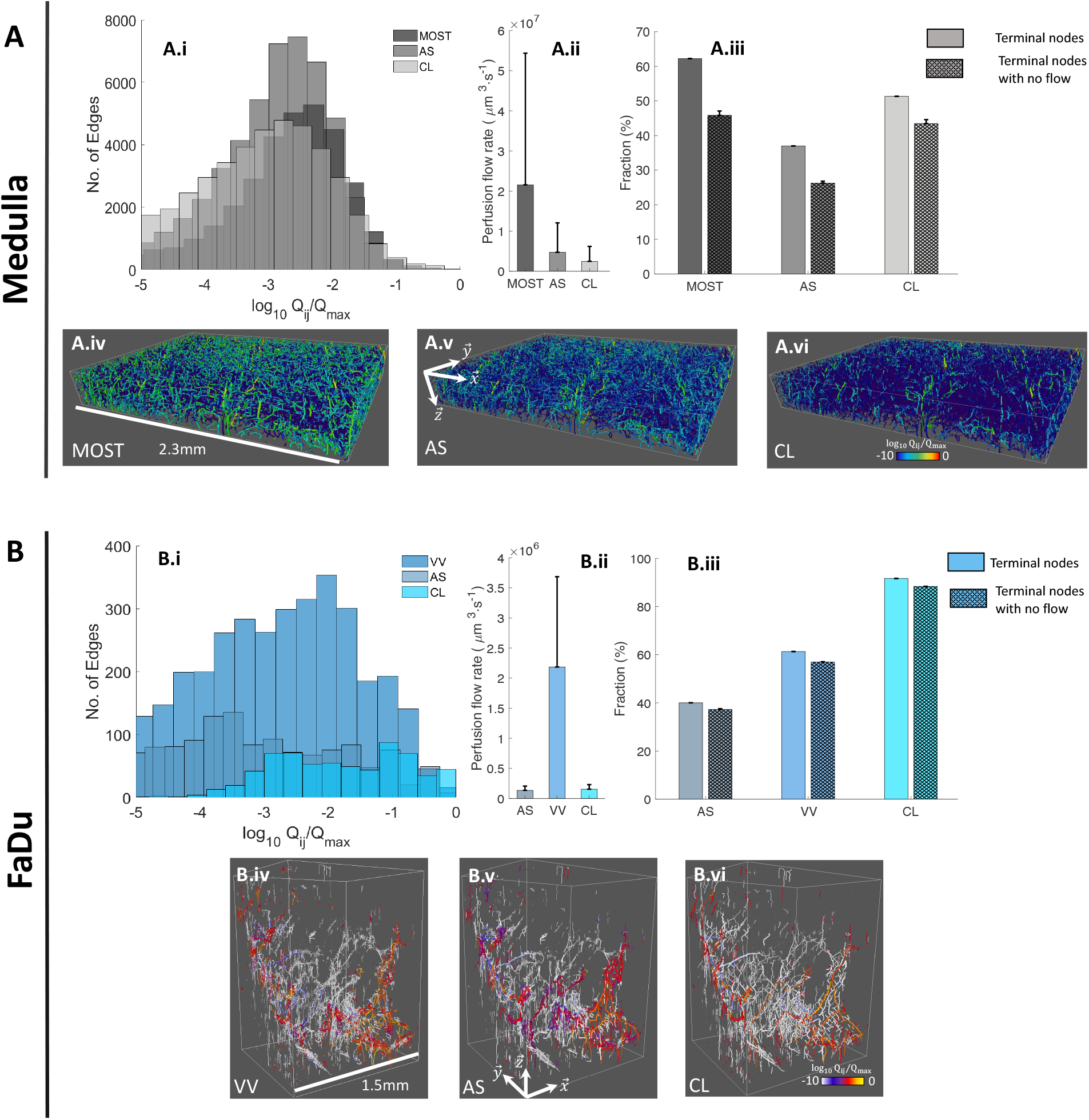
Flow simulation for (A) medulla and (B) FaDu networks for each skeletonisation algorithm. i) The flow distribution for the case of a pressure drop applied in the x direction. ii) The perfusion flow rate, i.e. the total flow rate going through the network, averaged over the three pressure drop directions, with the error bars representing the standard deviation. iii) Fraction of nodes that are terminal (plain), fraction of nodes that are terminal with no flow (cross hatch). iv) v) and vi) show the spatial flow distribution associated to the histogram displayed in i).

**Figure 7:**
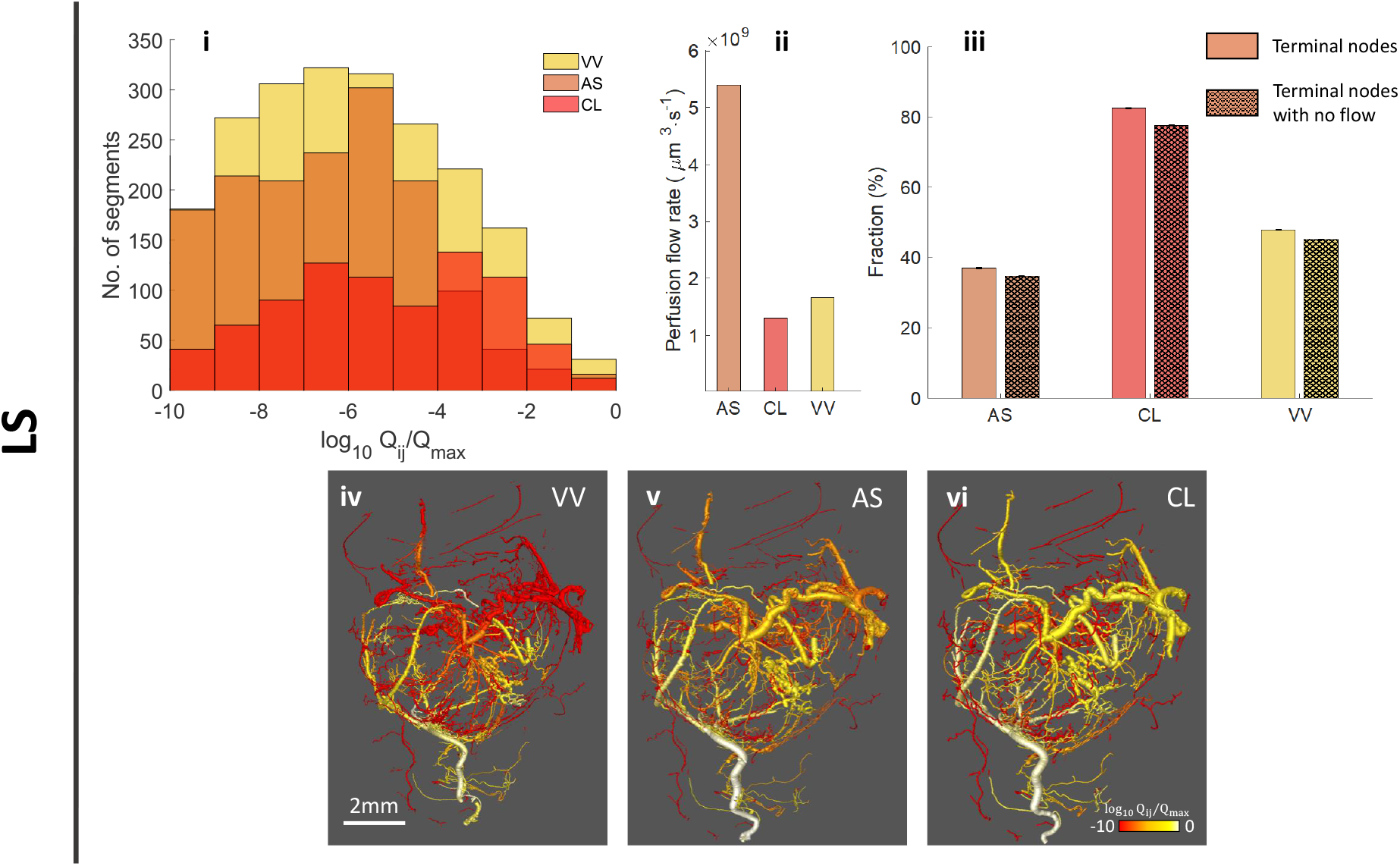
Flow simulation for the LS network. i) shows the flow distribution. ii) The perfusion flow rate, i.e., the total flow rate going through the network. iii) Fraction of nodes that are terminal (plain), fraction of nodes that are terminal with no flow (cross hatch). iv) v) and vi) show the spatial flow distribution associated to the histogram displayed in i).

Figures 6Ai, 6Bi and 7i show the flow rate distribution histograms for each network (for the case of a pressure drop along the 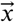 direction for the Medulla and FaDu networks). We can see that the distributions vary widely depending on the algorithm considered. Quantitatively, statistical comparisons (Table 3) show that flow distributions were significantly different across all networks, except the LS network in the case of CL against VV algorithms (Kruskall Wallis test with Dunn’s multiple comparisons, see details in Supplementary Table S10-S12). Such differences in flow distribution impact integral quantities such as the perfusion flow rate, defined as 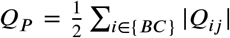 the total flow rate going through all terminal nodes (the set of terminal node being {*μC*}). Figures 6Aii and Bii, and 7ii show the resulting perfusion flow rate for the medulla, FaDu and LS networks respectively. We see that for each network there is consistently one skeletonisation algorithm associated with a perfusion flow rate approximately an order of magnitude larger than the others. We note, however, that there is no consistency across the three networks, with the medulla network showing largest perfusion flow rate when the MOST algorithm is used, the FaDU network when the VV algorithm is used and the LS network when the AS algorithm is used. Such differences also hold when looking at the net flow rates going through each face of the box-like shaped network (medulla and FaDu networks, see Supplementary Tables S7 and S8), regardless of the pressure drop direction considered. These differences in flow distribution and perfusion flow rates can be attributed to the cumulative effects of radius distribution, network connectivity and terminal node distribution.

As already mentioned in the Introduction and further illustrated by Equation (8), flow rates are particularly sensitive to the radius of blood vessels. Consequently, we see that the simulations predicting the largest perfusion flow rates are commonly associated with the algorithms reconstructing the network with the largest radii for the subsection networks (MOST algorithm for the medulla network (Figure 4Aii), VV algorithm the FaDu network (Figure 4Bii). For the LS network which is a complete network (as opposed to a box-like network) this pattern is broken; the AS algorithm despite having a lower mean radius that the VV algorithm produces the highest flow. This case highlights the complexity of linking network topology to dynamic behaviours, as can be seen from Figure 5iv there is a disconnect between one of the largest vessels in the LS networks produced by the VV algorithm that is not present in the CL or AS networks Fig. 5v and vi. This small change has a dramatic impact on the flow distribution in the skeleton produced by the VV algorithm, (Figure 5ii)).

Figures 6Aiv-Avii, 6Biv-Bvi, (medulla and FaDu networks, pressure drop along the 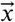 direction) and 7iv-vi (LS network) show strikingly different spatial flow patterns obtained depending on the algorithm used, reflective of the differences observed in flow rate distribution histograms. We see that in all cases the flow patterns are highly heterogeneous, which is a hallmark of the microvascular system (92). Still, we note the existence of regions associated with very small flow rates (dark blue for the medulla network, white for the FaDu network and red for the LS network), whose sizes vary depending on the algorithm considered. Such regions are the result of the combined effect of connectivity (number and spatial distribution of subnetworks) and boundary conditions (number and spatial distribution of terminal nodes). For instance, looking at the LS network and the flow prediction based on the VV algorithm (Figure 7iv), we see a large region with no flow on the top-right corner. We see that such a region is perfused with blood in the case of AS and CL algorithms (Figures 7v and 7vi). Recalling that we assigned the LS network with only one high pressure node (the node at the very bottom), i.e. one inlet, this means that this region, with relatively large segments, is actually considered by the VV algorithm as a disconnected, independent subnetwork.

Beyond highlighting the effect of radius distribution and network connectivity, the flow simulations demonstrate how different skeletonisation algorithms will lead to different terminal node distributions, and therefore to different boundary conditions. To illustrate this, Figures 6Aiii, Biii and 7iii show the number of terminal nodes, expressed as a fraction of the total number of nodes, in each network and for each algorithm (plain bars). We see that consistently at least 40% of nodes are terminal, increasing to 90% in the case of the CL algorithm and FaDu network. As already mentioned in section 4.2, the CL algorithm breaks looped structures, creating microvascular trees instead, which systematically increases the number of terminal nodes. Moreover, Figures 6Aiii, 6Biii and 7iii also show the number of terminal node that are associated with no flow, expressed as a fraction of the total number of nodes (cross-hatched bars), either because they are located far from the boundary or in a non perfusing, independent subnetwork. We see that this fraction generally follows closely the fraction of terminal nodes, indicating that a large number of terminal nodes are actually non perfusing, and therefore that most of the terminal nodes are not located close to the boundary (for the case of the medulla and FaDu network specifically).

Linear or weakly non-linear problems such as the blood flow problem described in Section 3.2.2 are mostly controlled by their initial and boundary conditions, which requires a degree of arbitrary decision whether motivated by physiology or pragmatism. Strategies have been developed to mitigate the arbitrary aspect of boundary condition assignment, such as periodic boundary condition for terminal nodes on the faces of box-shaped networks (10), or formulating the blood flow model (Equations (6)-(8)) as a constrained minimization problem (93), or even setting pressure or flow rate values on nodes associated with specific segments (e.g. arterioles, venules that are known entry/exit points for the network). However, such strategies have their limits; e.g., they do not solve the problem of dangling ends located far from the boundaries or disconnected subnetworks. An intuitive solution to solve that latter issue could be to either remove/discard small subnetworks such as isolated segments, or include new segments to artificially increase connectivity and obtain a single, all-connected network. However, dangling ends and disconnected subnetworks can be features of pathologies, e.g. where parts of the original healthy network have atrophied leaving isolated segments (10; 94) or where there is dis-regulated angiogenesis, e.g. caused by cancer cell angiogenic factor secretion (14; 6), which makes it challenging to alter arbitrarily the network structure. In this context it is critical to be able to quantify the uncertainties attached to a set of network skeletons, and be able to label their features as either being the result of a physiopathological processes or the result of a gap in the skeletonisation algorithms. Beyond this lies the propagation of such uncertainties to functional prediction. In particular, the perfusion flow rate underpins drug delivery (95; 6; 18); given the range of values displayed in Figures 6 and 7, it is challenging to make informative and quantitative predictions using such datasets. Similarly, net flow rates associated with faces for box-like shaped networks (Supplementary Tables S7 and S8) can be used to infer the local permeability tensor of the tissue associated with the reconstructed network. Such a permeability tensor, being a key player in multiscale models (84; 85), can easily propagate the uncertainties generated by the network skeleton to predictions at courser length scales.

In summary, the flow simulations highlight the limited ability of the skeletonisation algorithms to consistently recover key features of the flow, whether due to differences in the reconstructed segment radii, network connectivity or terminal node distribution. This motivates the need for a quantitative and easy to apply metric to assess and optimise skeletons produced by different algorithms.

## 5. A Metric for Skeleton validation

The creation of a full skeleton consensus ground-truth in an analogous manner to the segmentation fields is an overly manually intensive approach which could not be widely applied (24). An alternative approach is to consider that the skeletonization step should alter as little as possible the information contained in the segmented image. Doing so means to consider the segmented image as a gold standard, so that metrics derived from the skeleton then need to be compared to metrics derived from the segmented image. However, as revealed from our analysis in Section4.3 we should weight our comparison with the segmented image towards the information content of the segmented image that most affects dynamic behaviour. This includes: radius of vessels, connectivity of vessels in particular the connectivity of the largest single connected component of the networks which carries the majority of the blood flow (see Supplementary Figure S2). In addition to this, spatial encoding of geometric similarity must be included to ensure dynamic similarity, (note the principles of medialness and homology should ensure some spatial encoding of similarity). Such a comparison to the segmented image leverages the methods from the segmentation field used to create gold-standard segmentations, and provides a fixed point against which to assess a new skeletonised network in all cases. A number of other researchers have also sought to validate skeletons by comparison to binary images, (23; 5; 35), but these only consider one or two metrics in isolation, rather than formulating it as a collective. For example Table 4 shows comparison between the STAPLE segmentation and the skeletons for the medulla and FaDu Tumour datasets, across four different measures: Volume, number of connected components, Euler characteristic of the largest subnetwork (connected component), DICE score for bifircation points, and cl-sensitivity(33). It can be seen that depending on which metric is chosen for comparison, a different skeletonisation algorithm will appear to be superior. In the Medulla case, the CL algorithm gives the closest number of connected components to the STAPLE segmentation, however CL is the least good when the Euler number of the largest connected component or the bifurcations dice in a small subvolume is considered. The AS algorithm, on the other hand, provides the highest bifurcation DICE and cl-sensitivity, but has the lowest similarity in terms of volume and connected components compared to the other candidate skeletonization algorithms. These differences are also apparent in the FaDu data, but to an even greater extent owing to the pathology of this network and the lower resolution of the image data. This further highlights why a single metric which take into consideration multiple properties of a microvascular network is needed.

**Table 4.** Comparison of skeletonisations utilising individual metrics described above.

| Datasets | Volume<br>$\mu m^3$ | No. connected<br>components | Euler No.<br>of largest<br>connected component | Bifurcation DICE | cl- sensitivity |
| --- | --- | --- | --- | --- | --- |
| **Medulla** |  |  |  |  |  |
| **Binary Image** | **1.42** | **74** | **-381** | **1** | **1** |
| VV | 0.738 | 71 | -381 | 0.5 | 0.9733 |
| AS | 0.337 | 62 | -381 | 0.74 | 0.9999 |
| CL | 0.412 | 74 | 1 | 0.56 | 0.9997 |
| **FaDu** |  |  |  |  |  |
| **Binary Image** | **4.60** | **6067** | **-333** | **1** | **1** |
| VV | 3.80 | 3965 | -264 | 0.2 | 0.8997 |
| AS | 1.12 | 836 | -333 | 0.61 | 0.9499 |
| CL | 1.24 | 6067 | 1 | 0.35 | 0.9187 |

We have developed a composite metric that combines several morphological measurements of a skeletonized network and can be quickly computed from the network spatial graph. We term this the super-metric and it contains 5 measures each calculated by comparison to the binary image:

1. *V* - Total network volume
2. *μ* - Number and location of bifurcation points in a small subset of the network, reported as a DICE score
3. *cc* -The number of connected components or subgraphs of the network.
4. χ - The local Euler characteristic, where we have reformulated the classical Euler characteristic to resolve the special case of χ_*classical*_ = 0 . Our local Euler will always be positive χ = −χ_*classical*_ + 2. (35; 23; 96).
5. *cl* - A partial form of the cl-dice metric (only the sensitivity portion i.e. the overlap of the skeleton centreline with the binary image following) (33; 5).

For the *cl* measure we consider only the overlap of the centreline from the skeleton with the binary image, i.e the cl-sensitivity. *V, cc* and χ have equivalence in both the binary image and the spatial graph form. For *V, cc* the equivalence is trivial and for the χ number we compute the number of holes or tunnels through the largest connected component in the binary image, or equivalently, in the largest subgraph of the network, the number of nodes minus number of segments(96). Our variation from the classical Euler characteristic resolves the valid but special case where Euler characteristic would be zero, (e.g. a network with one single loop). For *μ* manual annotation of bifurcation points in a small subvolume of the binary data must be performed once, then the True positive, False Positive, and False negative bifurcation points in the same subregion can be computed for any skeleton automatically. *cl* is calculated by transformation of the spatial graph into a binary centreline image using Bresenham’s algorithm (97). The overlap between the two images is then used to calculate the images where one is the centreline of the spatial graph and the other the binary image. See supplementary Info section 7. and Supplementary Table 13 for calculation of each metric

These measures are combined the following way:

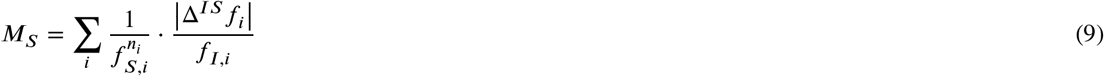

where *f*_*S,i*_ and *f*_*I,i*_ represent each of the metrics (as provided in Supplementary Table 13) computed from the reconstructed skeleton or the initial segmented image respectively. Then Δ^*IS*^*f*_*i*_ represent the difference in metric values between the two estimates. Finally, *n*_*i*_ are integers representing the weight given to each individual metrics. Taken together, we see that Equation 9 can be written as the projection (scalar product) of the distance vector between the reconstructed skeleton and segmented image onto a weighted space representative of the priority given to the different elementary metric.

The non-linear weighting for each term has been formulated to weight appropriately for the influence of each measure on the dynamic properties of the network. *V*, χ and *cc* all have *n* = 0, i.e. they are assigned with unit weights. On the other hand, the bifurcation DICE (*μ*) the cl-sensitivity (*cl*), elementary metric are weighted with *n* = 2 and *n* = 3 For the present case of the metric this gives a formulation:

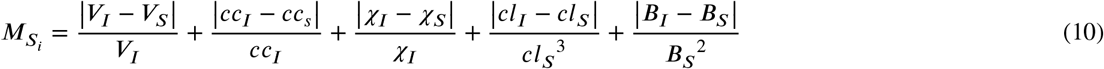

The practical affect of the weighting is that only a very minor reduction in *cl* from 1 will result in a high super-metric value, which is intuitively understandable i.e. a skeletonisation algorithm may occasionally cut a corner and still represent the network in the segmented image well, but large proportions of centreline outside of the binary segmentation is not plausible. For *μ*, larger deviation from 1 in the DICE bifurcation are tolerated by the super metric as the bifurcation points are annotated manually and hence may be slightly subjective. As *cl* or *μ* deviate from 1 the terms can quickly become dominant in the overall metric leading to a rejection of the skeleton, even if other morphological features are correct such as the number of connected components. This helps to enforce the spatial correctness which was highlighted as critical for dynamic behaviour.

In order to assess the performance of the super-metric, we performed formal optimisation of the four algorithms VV, CL, AS and MOST on a subsection of the medulla brain dataset See Supplementary information section 7 for optimisation methods.

Fig 8A shows that the super metric was able to provide a clear optimal algorithm choice from amongst the four algorithms-The AS algorithm had the lowest super metric score in the optimal case and indeed was generally lower than all other algorithms for most of the parameter spaces explored. Fig 8B shows how each term of the super metric contributes to the overall value of the super-metric for the baseline and optimal case for each algorithm. The very high DICE bifurcation case can be seen for the MOST algorithm indicating the spatial inaccuracy of this skeletonisation method. The super-metric also proves to be a highly useful tool in identifying, in which area an algorithm is performing most poorly. For example in Fig 8B it is clear that for the AS algorithm the Volume term is the highest contributor to the super-metric value. When we consider that this algorithm shows lower mean radius than its closest competitor the VV algorithm (also show in Supplementary Figure 5; it indicates that the method for evaluation of radius in the AS algorithm should be improved. Finally Figure8C shows how the absolute flow rate through the networks changes in response to optimisation of the super metric. As can be seen the optimal flow rates of the network are more tightly clustered as shown by the smaller 95% confidence intervals in the Optimal flow of Figure 8C.

**Figure 8:**
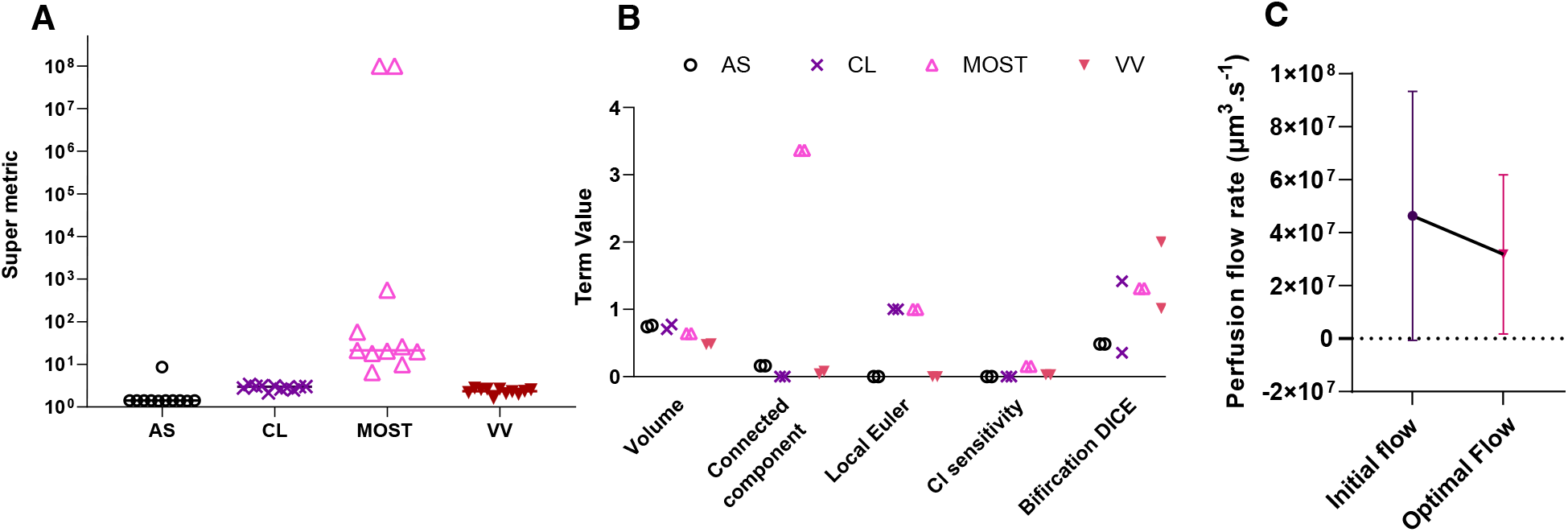
Showing how the supermetric optimisation impacts skeletons. **A)** the value of the supermetric in all 10 runs for each algorithm, **B)** each term in the super metric is plotted for the baseline, optimal and worst case of the parameter sets.**C)** The perfusion flow rate change i.e. the total flow rate going through the network between the initial and optimal parameter values for all four algorithms. The mean and 95% CI for all four networks in the initial and optimal case.

In summary, we have defined a super-metric for the assessment of skeletonisation algorithms. The super-metric can be considered a scalar product of a distance and weighting term for each component included. We have chosen to include 5 features in the term guided by our previous dynamic and structural metric study. We show that the super-metric can be easily and rapidly computed and applied to a wide range of skeletonisation algorithms. When this is done it is simple to compare different skeletonisation algorithms and choose that which provides the minimum super-metric value. In addition the individual terms of the super-metric can guide algorithm improvement. Whilst we believe this initial super-metric captures and combines the critical measures that should be used to assess micro-vascular network skeletons, the formulation is clearly amenable to being added to or adapted by others. In particular an addition that would allow radius and length to be distinguished (rather than a global volume term) would be preferable. Furthermore formulation of this metric in a numerically differentiable form could lead to its use as a loss function for machine-learning based skeletonisation approached.

## 6. Conclusion

The critical role of vascular networks in healthy tissue function and as biomarkers of disease is indisputable. This importance coupled to the increasing advances in 3D imaging techniques and computational frameworks for simulating the functional properties of networks from geometric data has huge potential in the field of computational medicine(20; 21). The image-processing pipeline is a key step in such image-based modelling frameworks of large microvascular networks. Ensuring that there is robust and widely agreed upon validation methods in place for the image processing portion of these frameworks is therefore crucial for their application in biological and medical scenarios.

In this work we have demonstrated the extent to which image processing variability, and in particular skeletonisation, can create variability in the structural properties measured from vascular networks and how this can propagate into frameworks that seek to model the functional properties of microvascular networks from 3D imaging data.

These differences can be so large that substantively different biological conclusions regarding e.g. the tortuosity or connectivity of any vascular network may be arrived at solely due to the selection of the skeletonisation algorithm even where it has parameterised by accepted methods in the field. This is critical where multiple studies are assessing how pathological changes in a vascular network may lead to functional differences in e.g. blood flow or oxygenation and what the biological implications of this on tissue function may be. The differences in functional properties (such as blood flow or perfusion) cannot be easily intuited from structural metrics owing to the complex dependencies on overall network connectivity, individual vessel properties such as radius, and how boundary conditions and nodes are identified and initialised.

Within the image processing pipeline the field of segmentation, has numerous widely applied metrics and strategies to mitigate variation caused by different algorithms or expert annotations. For skeletonisation the validation landscape is far less clear.

There are a wide variety of open source and commercial algorithms for performing skeletonisation with a vast and widely spread body of literature on the various developments and implementations. As a global goal, many of these algorithms have three guiding principles: homology preservation, thinness and medialness. Whilst these goals are, to a greater or lesser extent, achieved by individual algorithms, there is increasingly a push to scale such algorithms to handle the ever increasing size of 3D imaging data. With the added constraint of efficiency for data sets of 100GB and upwards, it may be that whilst the mathematical proof for e.g. homology holds in the original algorithm (50), efficient computational implementation through parallelisation, for example, produces unexpected behaviour or breaks assumptions of the original algorithm (52).

The fundamental challenge for skeletonisation, is that there is generally no gold-standard for a skeleton, or agreed upon metrics for assessing skeletons. The creation of a gold standard for skeletons is a large challenge. To the authors knowledge, the only published manually-defined consensus centreline datasets are the coronary artery datasets of Schaap et al. (24), where the length of time required to produced consensus centrelines for just the first 3 branches of the coronary artery tree was over 500 hours of expert time. Instead we have proposed a super-metric for skeletonisation that utilised the binary input image as the gold-standard and seeks for the skeleton to most closely represent the binary image with a weighting to features which most heavily affect the functional behaviour of the network. To that end we have proposed a metric which consists of 5 terms each of which captures features that we have demonstrated are important to the functional behaviour of microvascular networks, and we have combined these as a scalar product of a distance from the binary image and a weighting term. This formulation is simple and fast to compute and thus is practically usable by any researcher as well as providing intuitive insights into the ways in which a particular skeletonisation algorithm differs from other algorithms or from the binary input image used to create it. We aim that this super-metric become a standard metric for assessing microvascular skeletons as it will allow of a quantitative assessment of new algorithms and will empower researchers to be able to effectively choose and optimise new skeletonisation algorithms. Further development of the super-metric with the addition of new terms of the refinement of our existing terms would be highly beneficial particularly in assessing how the radius of vessel is estimated, or by efficiently including functional properties in the metric composition.

## Abbreviation Term

χ: Local Euler characteristic
χ_*classical*_: Classical Euler characteristic
*μ*: blood effective viscosity
*τ*_*ij*_: Tortuosity of a segment
*θ*: Branching angle
AS: Auto Skeleton
*μ*: Bifircation point DICE score
BC: Set of terminal nodes at which boundary conditions will be set
*cc*: Number of connected components
CL: Centreline Tree
*cl*: Centreline sensitivity
CT: Computed Tomography
DTHO: Distance Transformed Homotopic Ordering FN False negative
FP: False positive
*G*_*s*_: conductance of subsegment
*H*: Hausdorff distance
*H*_*d*_: Haematocrit
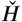: Directed Hausdorff fistance
HiP-CT: Hierarchal Phase Contrast Microcscopy ICM Inflection count metric
IVD: Intervessel distance
*L*_*s*_: Length sub-segment
LDR: Length to diameter ratio
LS: LS174T
*M*_*S*_: Super metric for algortihm S
MOST: Micro Optical sectioning tomography
MR-HREM: Multi-fluorescent High Resolution Episcopic Microscopy MRI Magnetic resonance imaging
OPT: Optical projection tomography
*P*_*i*_: Pressure at node i
*Q*_*s*_: Flow rate in a subsegment
*R*_*s*_: Radius sub-segment
*s*: Sub-segment
SOAM: Sum of angle metric
STAPLE: Simultaneous truth and performance level estimation TEASAR Tree structure extraction algorithm
TP: True positive
*V*: Volume
*V*_*c*_: Volume consensus segmentation
*V*_*t*_: Volume of test segmentation
VV: VesselVio

## Acknowledgements

This work has received funding from, EPSRC (Grant nos. EP/R004463/1 and EP/W007096/1), the Imaging-BioPro Networks BBSRC/EPSRC/MRC (grant no. MR/R0256731/1), Cancer Research UK(C44767/A29458 and C23017/A27935), the Chan Zuckerberg Initiative DAF, an advised fund of the Silicon Valley Community Foundation, grant number 2020-225394 and an MRC Skills Development fellowship (MR/S007687/1), The Rosetrees Trust (AN18546).

