## Supplementary information for "Reconstructing microvascular network skeletons from 3D images: what is the ground truth?"

### Reconstructing micro-vascular networks, from 3D images: where is the ground truth?
### Supplementary Information

### 1 Dataset parameters

Parameter definition for each Skeletonisation method

- Centerline Tree (slope, zeroVal,Number of Parts),
- Auto skeleton (smooth y/n, smooth, attach to data, iterations)
- MOST (Threshold, Seed_size, slip_size, Pruning (y/n))
- VesselVio(filtering isolated segments y/n, pruning y/n, anisotropic y/n)

| Spatial graph filename | Imaging modality | Sample | Resolution of images (um) | Segmentation method | Skeletonisation method with params |
|---|---|---|---|---|---|
| Medulla_centreline | MF-HREM | BALB/C Mouse 12 weeks brain medulla region HREM | 1.14 x 1.14 x 1.72 | STAPLE of all manuals | Centreline tree (2.5,4,-1) |
| Medulla_MOST | | | | MOST | MOST (5,6,10, n) |
| Medulla_autoskeleton | | | | STAPLE of all manuals | Auto skeleton (n,na,na,na) |
| FaDu_centeline | MF-HREM | BALB/c 12 weeks FADU tumour left flank | 2.75 x 2.75 x 2.58 | STAPLE all manuals | Centreline tree (2.5,4,-1) |
| FaDu_autoskeleton | | | | STAPLE all manuals | Auto skeleton (n,na,na,na) |
| FaDu_VesselVio | | | | STAPLE all manuals | Vessel Vio (n,n,y) |
| LS_centreline_tree | CT | BALB/c 12 weeks LS tumour | 22 x 22 x 22 | STAPLE all manuals | Centreline tree (2.5,4,-1) |
| LS_autoskeleton | | | | STAPLE all manuals | Auto skeleton (n,na,na,na) ) |
| LS_VesselVio | | | | STAPLE all manuals | Vessel Vio (n,n,n) |

**Table S1.** parameters for each dataset for segmentation and skeletonization**.**

The datasets for these three samples can be found from Walsh et al. 2021, and Holroyd et al. 2023

#### 1.1 Parameter selection for the data set Medulla_MOST

The parameter selection process was done after Wu et al. 2014, in brief, image slices were extracted every 12th slice from the top of the STAPLE segmented image volume. The vessel cross sections in these slices were considered ground truth and each connected component in a slice is termed a region. The graph output of the MOST pipeline was converted into a binary image using Vaa3D, and every 12th slice used for comparison to the ground truth. A vessel was denoted as correctly traced if there was overlap between the binary conversion of the graph file and the ground truth slice region. The recall, denoted by $R_b$, was defined as $R_b = B/B_1$where $B$ is the number of overlap regions and $B_1$is the total number of ground truth regions. The precision, denoted by$P_b$, was defined as $P_b = B/B_2$ where $B_2$ represents the total number of traced regions.

We combined the precision and recall into a single F1 score for ease of comparison as:

$$F1 = 2\frac{P_b * R_b}{P_b + R_b}$$

Using this approach the highest F1 score was used to select the MOST parameters. This wa found to be 5,6, and 20 for the threshold, seed and slip or scoop[1] respectively with an F1 of 0.83.

| Threshold | Slip or Scoop | Seed | F1 |
|---|---|---|---|
| 20 | 6 | 20 | 0.48 |
| 3 | 4 | 20 | 0.81 |
| 43 | 6 | 20 | 0.24 |
| 5 | 4 | 20 | 0.8 |
| 5 | 6 | 10 | 0.83 |
| 5 | 6 | 5 | 0.83 |
| 8 | 2 | 20 | 0.71 |
| 8 | 6 | 20 | 0.71 |

**Table S2.** parameters search for MOST pipeline. Red indicates the highest F1 score, Slip or Scoop is the radios for the scooping process, at each iteration and seed is the size (voxels) of the initial seed point

[1] This parameter is the scoop parameter from Rodriguez et al. 2009 but is called slip in the Vaa3D implementation of the algorithm.

#### 2 Spatial Graph Definition and Metrics

This section shows how a spatial graph is defined in terms of nodes, segments, points and subsegments, and shows how structural metrics are calculated from this representation.

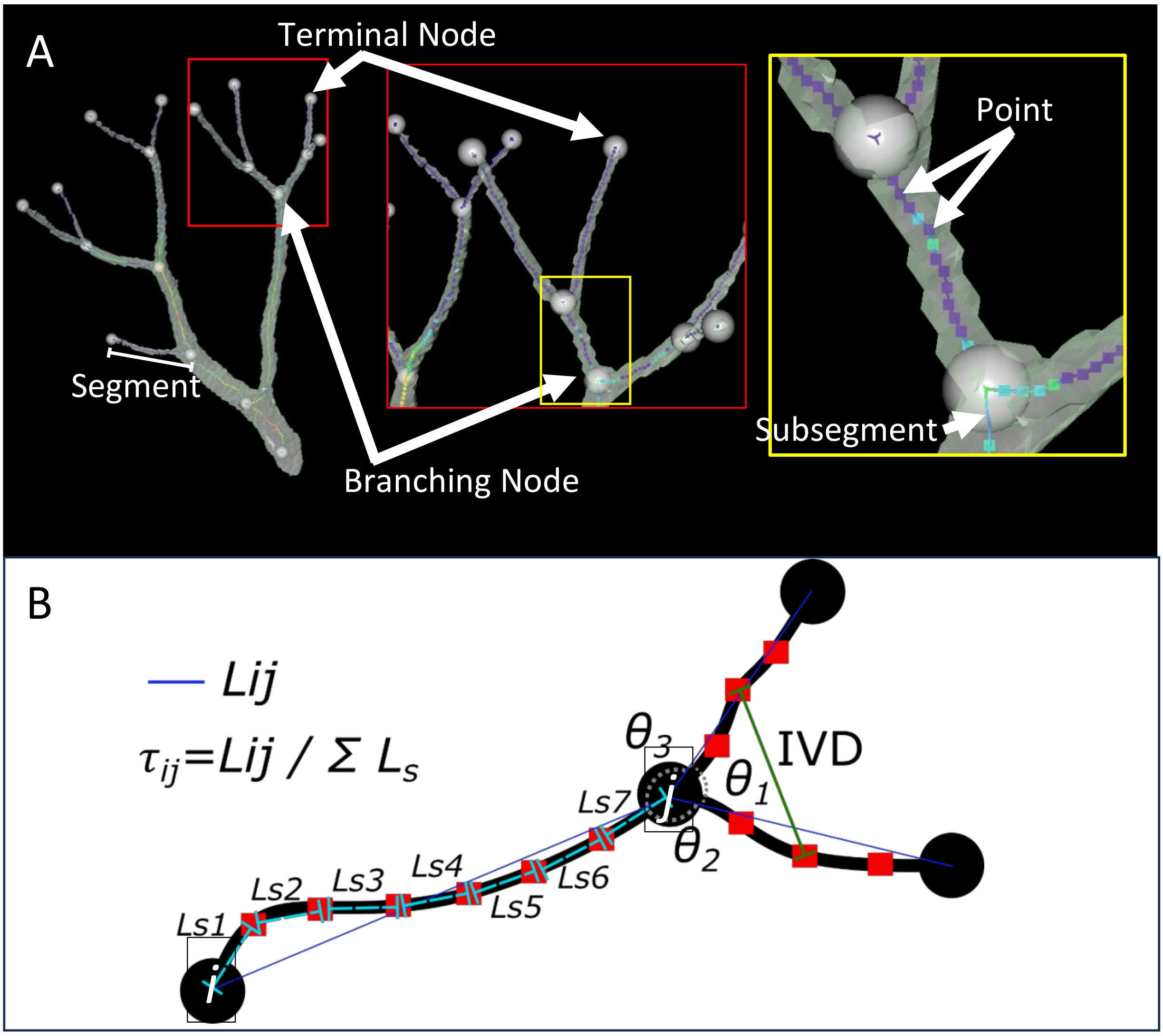

**Figure S1.** Figure showing A) how the skeletonised is defined relative to the segmented object in terms of terminal nodes, branching nodes, segments, subsegments and points. B) How the metrics are calculated from the skeletonised graph networks, where for every segment is describe by start node *i*, end node *j and subsegments s. The tortuosity* $\tau$, *and branching angles* $\theta_1, \theta_2, \theta_3$ *are shown.*

### 3 Spatial graph Summary statistics

| Spatial graph | Number Nodes | No. subgraphs | No. segments | Total volume ($\mu m^3$) | No. Branched nodes | No. terminal nodes |
|---|---|---|---|---|---|---|
| **Medulla Centreline Tree** (CL) | 154267 | 1648 | 152619 | $9.1 \times 10^7$ | 75093 | 79174 |
| **Medulla MOST** | 94726 | 9792 | 84667 | $1.2 \times 10^8$ | 35476 | 58944 |
| **Medulla Auto Skeleton** (AS) | 58006 | 1310 | 66948 | $8.86 \times 10^7$ | 36578 | 21428 |
| **FADUS Centreline Tree** (CL) | 14575 | 6067 | 8508 | $1.24 \times 10^7$ | 1212 | 13363 |
| **FADUS Auto Skeleton** (AS) | 6330 | 836 | 7959 | $1.12 \times 10^7$ | 3979 | 2351 |
| **FADUS VesselVio** (VV) | 19635 | 3965 | 18056 | $3.8 \times 10^7$ | 7681 | 11951 |
| **LS Centreline Tree** (CL) | 3704 | 1202 | 2502 | $5.60 \times 10^9$ | 647 | 3057 |
| **LS Auto Skeleton** (AS) | 4378 | 339 | 4378 | $5.31 \times 10^9$ | 2028 | 1067 |
| **LS VesselVio** (VV) | 6348 | 812 | 7041 | $1.15 \times 10^{10}$ | 3366 | 2981 |

**Table S3.** Summary of graphs tolopological characteristics of each network (Medulla, FaDu, LS) and each skeletonization algorithm (AS, CL, VV, MOST).

| Spatial graph | No. of subgraphs of >10% of total network volume |
|---|---|
| **Medulla Centreline Tree** (CL) | 1 |
| **Medulla MOST** | 1 |
| **Medulla Auto Skeleton** (AS) | 1 |
| **FADUS Centreline Tree** (CL) | 3 |
| **FADUS Auto Skeleton** (AS) | 4 |
| **FADUS VesselVio** (VV) | 2 |
| **LS Centreline Tree** (CL) | 2 |
| **LS Auto Skeleton** (AS) | 2 |
| **LS VesselVio** (VV) | 2 |

**Table S4.** Subgraph volume distribution

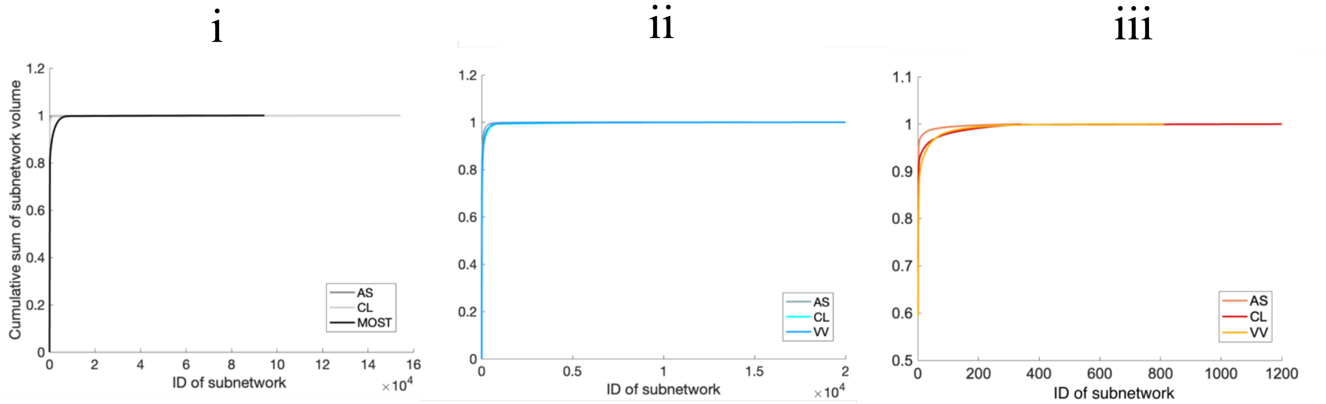

**Figure S2.** Cumulative sum of the volume for subnetworks for each network i)Medulla, ii) FaDu, iii) LS. Each skeletonization algorithm (Auto Skeleton (AS), Centreline Tree (CL), VesselVio (VV), and MOST).

Figure S3. And Table S4 both show the volume proportion for each subnetwork of the total spatial graph. In Table S4 the total number of subnetworks with more than 10% of the total network volume are presented, in all cases this is 4 or less, and in majority of cases it is 2 or less. This feature whereby, most of the volume of the total network is contained within only a few of the subnetworks is demonstrated graphically in Figure S3. Here we rank and then plot the normalised cumulative volume for each subnetwork, again it can be seen that in all cases the majority of the network volume is contained within the first 1-5 subnetworks.

| **Sample** | **Volume μm$^3$** | **No. connected components** |
|---|---|---|
| **Medulla** STAPLE | $3.2 \times 10^8$ | 1648 |
| **FaDu** STAPLE | $4.6 \times 10^7$ | 6067 |
| **LS** STAPLE | $1.24 \times 10^{10}$ | 1202 |

**Table S5.** Volume and number if connected components (26 neighbourhood) for the binary image volumes from the consensus segmentation.

### 4 Boundary Conditions for flow simulation

Two approaches for boundary conditions are applied for box-shaped networks (i.e. FaDu and Medulla, Figure S4) or for complete network (i.e. LS tumour, Figure S5).

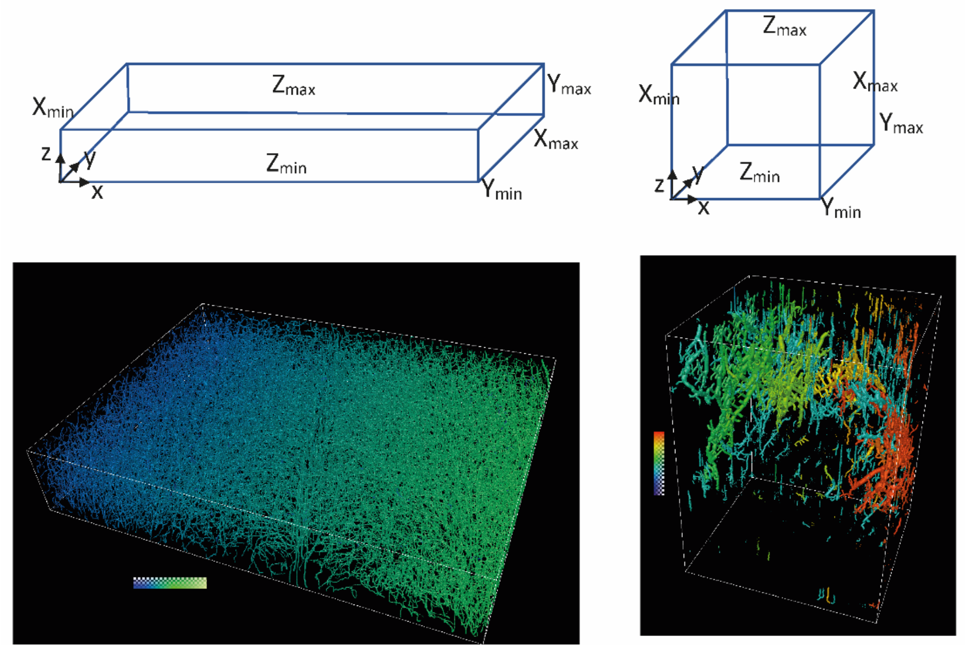

**Figure S3.** pressure boundary condition examples for partial networks of medulla and FADU. Both networks show the case where the pressure gradient is imposed at the Xmin and Xmax faces.

| Centreline Tree (CL) | | Auto Skeleton (AS) | | VesselVio (VV) | |
|---|---|---|---|---|---|
| total node num 3704 | | total node num 3095 | | total node number 6348 | |
| **Node number*** | **subgraph** | **Node number*** | **subgraph** | **Node number*** | **subgraph** |
| 57 | 0 | 3091 | 0 | 5925 | 0 |
| 53 | 0 | 1487 | 0 | 2911 | 2 |
| 23 | 0 | 820 | 0 | 1895 | 4 |
| 1044 | 1 | 2403 | 1 | 4661 | 1 |
| 2740 | 2 | 3021 | 2 | 5843 | 3 |
| 1691 | 3 | 1170 | 4 | 3237 | 6 |
| 2638 | 4 | 1992 | 3 | 3908 | 5 |
| *Numbering starting from 1 | | | | | |

**Table S6.** boundary nodes shared across LS skeletons

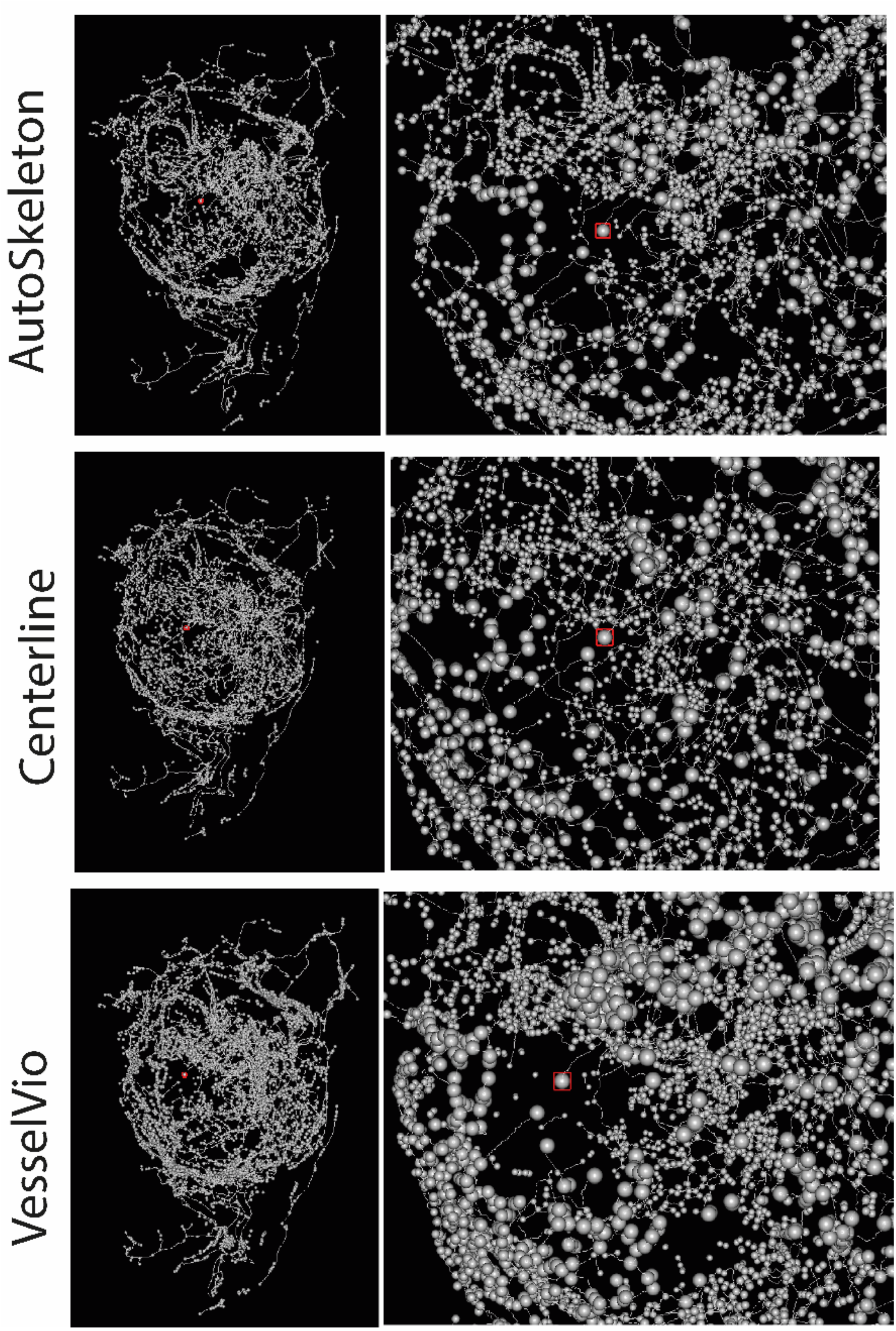

**Figure S4.** Illustration of the node chosen (highlighted in red) manually for each skeleton of the LS network.

### 5 Results of flow simulations

| Skeletonisation Algorithm | BC imposed | Xmin (µm³/s) | Xmax (µm³/s) | Ymin (µm³/s) | Ymax (µm³/s) | Zmin (µm³/s) | Zmax (µm³/s) |
|---|---|---|---|---|---|---|---|
| **Auto Skeleton** (AS) | Pressure drop across x | 6.1E+03 | -1.8492.3 | -4.7E+03 | 2.0E+03 | 1.0E+04 | 4.8E+03 |
| | Pressure drop across y | 2.6E+02 | -9.1E+02 | 1.8E+04 | -3.1E+04 | 2.8E+05 | -2.6E+05 |
| | Pressure drop across z | -2.4E+04 | -1.4E+05 | 4.4E+05 | -1.6E+05 | 6.0E+06 | -6.2E+06 |
| **Centreline Tree** (CL) | Pressure drop across x | 2.9E+03 | -7.6E+03 | -5.8E+03 | 8.6E+02 | 1.2E+04 | -2.3E+03 |
| | Pressure drop across y | -1.0E+03 | -2.8E+03 | 7.9E+03 | -1.2E+04 | 1.9E+05 | -1.9E+05 |
| | Pressure drop across z | -6.1E+04 | -1.2E+05 | 3.5E+04 | -8.0E+04 | 3.7E+06 | -3.4E+06 |
| **MOST** | Pressure drop across x | 2.6E+04 | -5.1E+04 | -6.6E+03 | -3.1E+03 | 4.2E+04 | -6.7E+03 |
| | Pressure drop across y | 2.3E+03 | -6.8E+03 | 6.7E+04 | -1.3E+05 | 5.0E+05 | -4.3E+05 |
| | Pressure drop across z | -4.4E+04 | -5.1E+05 | 3.8E+05 | -6.2E+05 | 1.2E+07 | -1.1E+07 |

**Table S7.** Net flow rates perfusing each face (Xmin, Xmax, Ymin, Ymax) of the Medulla network.

| Skeletonisation Algorithm | BC imposed | Xmin (µm³/s) | Xmax (µm³/s) | Ymin (µm³/s) | Ymax (µm³/s) | Zmin (µm³/s) | Zmax (µm³/s) |
|---|---|---|---|---|---|---|---|
| **Auto Skeleton** (AS) | Pressure drop across x | 2.7E+02 | -2.3E+02 | 2.8E+02 | 1.3E+04 | -1.3E+04 | -2.4E+02 |
| | Pressure drop across y | 1.0E+02 | -4.1E+01 | -3.2E+01 | -3.4E+04 | 3.4E+04 | 2.8E+01 |
| | Pressure drop across z | -6.8E+02 | -1.6E+01 | 3.2E+02 | -4.0E+04 | 4.0E+04 | -3.1E+02 |
| **Centreline Tree** (CL) | Pressure drop across x | 5.8E+03 | -2.6E+03 | 1.7E+03 | 1.7E+04 | -2.1E+04 | -1.5E+03 |
| | Pressure drop across y | -4.4E+02 | 2.1E+04 | 2.3E+02 | -5.5E+04 | 3.4E+04 | -2.7E+02 |
| | Pressure drop across z | -7.8E+03 | 2.0E+04 | -1.4E+03 | -5.6E+04 | 4.5E+04 | 3.7E+02 |
| **VesselVio** (VV) | Pressure drop across x | 1.4E+04 | -4.2E+02 | -4.2E+03 | 8.0E+04 | -9.4E+04 | 4.6E+03 |
| | Pressure drop across y | -4.0E+03 | 6.5E+04 | 1.4E+03 | -2.3E+05 | 1.6E+05 | 3.7E+03 |
| | Pressure drop across z | -4.7E+04 | 2.3E+04 | -4.7E+03 | -2.2E+05 | 2.6E+05 | -1.6E+04 |

**Table S8.** Net flow rates perfusing each face (Xmin, Xmax, Ymin, Ymax) of the FaDu tumour network.

| Skeletonisation Algorithm | Total flow (µm³/s) |
|---|---|
| **Auto Skeleton** (AS) | 5.4E+08 |
| **Centreline Tree** (CL) | 1.3E+08 |
| **VesselVio** (VV) | 1.6E+09 |

**Table S9.** Total flow perfusing the LS tumour network.

#### 6 Output of statistical tests

| **Skeletonisation algorithm pair** | **Radius** Df (2) $\chi^2(47829.4)$ p (0) | **Branching** Df (2) $\chi^2(184.832)$ p (0) | **IVD** Df (2) $\chi^2(1517)$ p (0) | **Tortuosity** Df (2) $\chi^2(61960)$ p (0) | **LDR** Df(2) $\chi^2(20220.3)$ p (0) | **Flow** Df (2) $\chi^2(38112)$ p <0.0001 |
|---|---|---|---|---|---|---|
| **MOST** vs **Auto Skeleton** (AS) | 0 | 3.33E-16 | 0 | 0 | 0 | <0.0001 |
| **MOST** vs. **Centreline Tree** (CL) | 0 | 0.000611 | 0 | 0 | 0 | <0.0001 |
| **Auto Skeleton** (AS) vs **Centreline Tree** (CL) | 0 | 0 | 0 | 0 | 0 | <0.0001 |

**Table S10.** Statistical output for structural and functional metric distribution, Kruskall Wallis test with Dunn's correction for multiple comparison for the Medulla network.

| **Skeletonisation algorithm pair** | **Radius** Df (2) $\chi^2(8637)$ p (0) | **Branching** Df (2) $\chi^2(16.913)$ p (0.000213) | **IVD** Df (2) $\chi^2(4106.2)$ p (0) | **Tortuosity** Df (2) $\chi^2(3091.7)$ p (0) | **LDR** Df(2) $\chi^2(5792.23)$ p (0) | **Flow** Df (2) $\chi^2(593.3)$ p <0.0001 |
|---|---|---|---|---|---|---|
| **VesselVio** (VV) vs **Auto Skeleton** (AS) | 0 | 0.66874 | 0 | 0.00194 | 0 | <0.0001 |
| **VesselVio** (VV) vs. **Centreline Tree** (CL) | 0 | 0.000659 | 0 | 0 | 0.000109 | <0.0001 |
| **Autoskeleton** (AS) vs **Centreline Tree** (CL) | 0 | 0.000255 | 0 | 0 | 0 | <0.0001 |

**Table S11.** Statistical output for structural and functional metric distribution, Kruskall Wallis test with Dunn's correction for multiple comparison for the FaDu network.

| **Skeletonisation algorithm pair** | **Radius** Df (2) $\chi^2(2021.2)$ p (0) | **Branching** Df (2) $\chi^2(4.126)$ p (0.127) | **IVD** Df (2) $\chi^2(3638.2)$ p (0) | **Tortuosity** Df (2) $\chi^2(218.6)$ p ($3.47 \times 10^{-48}$) | **LDR** Df(2) $\chi^2(1941.4$[illegible] p (0) | **Flow** Df (2) $\chi^2(476.3)$ p <0.0001 |
|---|---|---|---|---|---|---|
| **VesselVio** (VV) vs **Auto Skeleton** (AS) | 0 | 0.608 | 0 | 0.0055 | 0 | <0.0001 |
| **VesselVio** (VV) vs. **Centreline Tree** (CL) | 0 | 0.77 | 0 | 0 | 0 | 0.2413 |
| **Autoskeleton** (AS) vs. **Centreline Tree** (CL) | 0.26141 | 0.131 | 0 | 0 | 0 | <0.0001 |

**Table S12.** Statistical output for structural and functional metric distribution, Kruskall Wallis test with Dunn's correction for multiple comparison for the LS network.

### 7 Proposed metric and optimization

We propose a skeletonisation meta-metric (which combines several morphological measurements of a skeletonized network) and can be compared these to the original binary segmentation from which the skeleton was produced. The super metric contains 5 components each a comparison to the binary segmentation.

1. Total network volume $V$
2. Euler number of the largest connected component $\chi = N - E$
3. Number of connected components $cc$
4. The DICE score for the branching nodes in randomized sub volume of the network. $B$ (range 0-1)
5. Cl-sensitivity score[2]. $Cl$ (range 0-1)

| Measure | Explanation | Calculation from binary image | Calculation from spatial graph |
|---|---|---|---|
| **V (Volume)** | Total network volume | Sum of all labelled voxels | Sum of volume of all subsegments |
| **B (Bifurcation DICE)** | The DICE score for each bifurcation point in a small subset of the data | Manually annotated in a subregion of the image | Check for each node in the spatial graph subregion for proximity to the 3D location in the manual annotation (a tolerance of proximity can be set) |
| **cc (no. of connected components)** | The number of unconnected regions | Number of connected components (26 neighbourhood) | Number of sub networks |
| **χ (Local Euler characterisitic)** | We have reformulated the classical Euler characteristic to resolve the special case of $\chi_{classical} = 0$. Our local Euler will always be positive $\chi = -\chi_{classical} - 2$ | For the largest connected component, the number of tunnels or holes within it | For the largest subgraph no. of nodes - no. of segments (Youssef 2015, Brown 2022, Chang 2021) |
| **cl (cl-sensitivity)** | A partial form of the cl-dice metric (only the sensitivity portion i.e. the overlap of the skeleton centreline with the binary image following (Shit et al. 2021) | NA | Transformation to a binary image of lines ($l_s$) via Bresenham algorithm (Bresenham et al. 1998) then: $\sum(V_I \cdot l_s)/\sum l_s$ |

**Table S13.** Each term in the super metric and how it is calculated in the binary image and spatial graph.

[2] shit2021cldice

Given that the four skeletonisation algorithms were all included in different library/software that were hardly scriptable, optimizing was manual intensive. To mitigate this effect we built, for each algorithm, a surrogate model using standard universal Kriging with spherical variogram model and looked for the optimal parameter combinations directly using the surrogate model (Virdee et al 1984, Murphy et al 2014). Precisely, we sampled the parameter space of each algorithm 10 times using Latin-Hypercube sample, which ensure adequate spread across the parameter space. For each algorithm the variable parameters are show below in bold.

- Centerline Tree (**slope**, **zeroVal**, Number of Parts),
- Auto skeleton (smooth y/n, **smooth, attach to data, iterations**)
- MOST (**Threshold, Seed_size, slip_size**,)
- VesselVio (**filtering isolated segments (length), pruning (length**), voxel size)

The parameter ranges used in the Latin-Hypercube sample were taken from hard limits or from reasonable limits given the size of the vascular features of interest.

| **Algorithm** | **Parameter** | **Min** | **Max** |
|---|---|---|---|
| **CL** | Slope (multiplier for dist value) | 1 | 6 |
| **CL** | zeroVal (pxl included as boarder) | 1 | 10 |
| **CL** | Number parts | -1 | -1 |
| **AS** | smooth | 0 | 1 |
| **AS** | Attach to data | 0 | 1 |
| **AS** | iterations | 1 | 15 |
| **VesselVio** | Isolated segment filter | 0pxl | 10pxl (14.4um) |
| **VesselVio** | Pruning length | 0pxl | 5pxl (5.7um) |
| **MOST** | Threshold | 1 | 40 |
| **MOST** | Seed (size pixel seed) | 2 | 10 |
| **MOST** | Slip (movement in pxl at each iteration) | 2 | 10 |

**Table S14.** Parameter limits for each algorithm as input to the Latin Hyperparameter sampling

Following the runs super-metric performance was calculated for each run, these values were used to create an analytical surrogate function which could be optimized. In each case the 0$^{th}$ run is the baseline used in the previous analysis.

| NAME | THRESHOLD | SEED | SLIP | SUPER- METRIC |
|---|---|---|---|---|
| MOST_0 | 5 | 6 | 10 | 6.48 |
| MOST_1 | 10 | 8 | 3 | 26.08 |
| MOST_2 | 22 | 9 | 4 | 19.97 |
| MOST_3 | 16 | 5 | 8 | 20.65 |
| MOST_4 | 27 | 7 | 9 | 554.75 |
| MOST_5 | 35 | 3 | 10 | 21.43 |
| MOST_6 | 7 | 10 | 3 | 9.85 |
| MOST_7 | 38 | 4 | 7 | $1x10^8$ |
| MOST_8 | 19 | 5 | 2 | 18.12 |
| MOST_9 | 2 | 8 | 8 | 57.72 |
| MOST_10 | 29 | 2 | 6 | $1x10^8$ |

**Table S15.** Parameters for MOST with output of super metric. Blue test indicates the baseline measure and green indicates the optimal parameter values.

| RUN_NAME | ISOLATED SEGMENT FILTER | PRUNING LENGTH | SUPER- METRIC |
|---|---|---|---|
| VV_0 | 0 | 0.00 | 2.55 |
| VV_1 | 5.00 | 1.57 | 2.62 |
| VV_2 | 2.07 | 5.49 | 1.61 |
| VV_3 | 12.56 | 4.65 | 2.07 |
| VV_4 | 7.98 | 4.35 | 1.99 |
| VV_5 | 3.49 | 1.03 | 2.62 |
| VV_6 | 5.89 | 3.12 | 2.30 |
| VV_7 | 1.43 | 1.97 | 2.58 |
| VV_8 | 10.19 | 2.63 | 2.35 |
| VV_9 | 9.78 | 3.69 | 2.19 |
| VV_10 | 13.83 | 0.31 | 2.78 |

**Table S16.** Parameters for VV with output of super metric. Blue test indicates the baseline measure and green indicates the optimal parameter values.

| RUN_NAME | SLOPE | ZEROVAL | SUPER-METRIC |
|---|---|---|---|
| CL_0 | 2.5 | 4 | 3.13 |
| CL_1 | 5.81 | 2.6 | 2.13 |
| CL_2 | 4.95 | 9.2 | 3.00 |
| CL _3 | 3.94 | 4.8 | 2.64 |
| CL _4 | 1.22 | 7.6 | 3.05 |
| CL _5 | 2.91 | 6.4 | 2.78 |
| CL _6 | 5.28 | 1.2 | 2.82 |
| CL _7 | 2.09 | 2.9 | 3.39 |
| CL _8 | 1.71 | 4.6 | 3.15 |
| CL _9 | 4.03 | 8.3 | 2.99 |
| CL _10 | 3.37 | 5.8 | 2.52 |

**Table S17.** Parameters for CL with output of super metric. Blue test indicates the baseline measure and green indicates the optimal parameter values.

| RUN_NAME | SMOOTH | ATTACH TO DATA | ITERATIONS | SUPER-METRIC |
|---|---|---|---|---|
| AS_0 | 0.5 | 0.25 | 10 | 1.41 |
| AS _1 | 0.06 | 0.07 | 5 | 1.39 |
| AS _2 | 0.96 | 0.78 | 3 | 1.42 |
| AS _3 | 0.83 | 0.62 | 14 | 1.42 |
| AS _4 | 0.57 | 0.41 | 7 | 1.42 |
| AS _5 | 0.27 | 0.86 | 10 | 1.42 |
| AS _6 | 0.66 | 0.35 | 12 | 1.41 |
| AS _7 | 0.75 | 0.51 | 4 | 1.42 |
| AS _8 | 0.35 | 0.95 | 8 | 8.67 |
| AS _9 | 0.43 | 0.12 | 2 | 1.39 |
| AS _10 | 0.10 | 0.26 | 12 | 1.41 |

**Table S18.** Parameters for AS with output of super metric. Blue test indicates the baseline measure and green indicates the optimal parameter values.

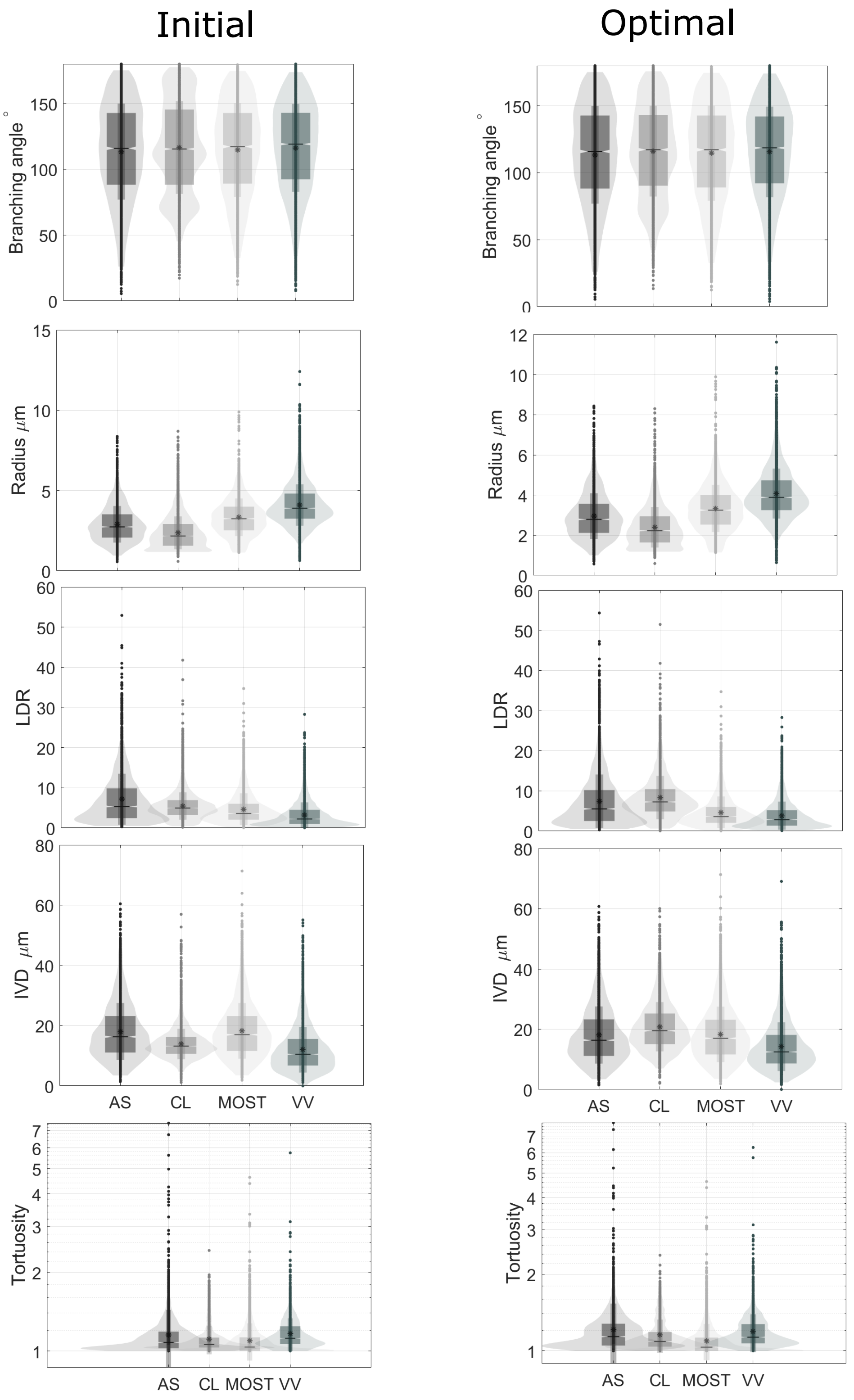

**Figure S5. The structural metrics for the initial and optimal parameter values of the medulla brain subsection**
